# HiChIA-Rep quantifies the similarity between enrichment-based chromatin interactions datasets

**DOI:** 10.64898/2026.02.01.703086

**Authors:** Sion Kim, Joseph T. Jackson, Henry B. Zhang, Minji Kim

## Abstract

**Background:** Enrichment-based 3D genome mapping technologies ChIA-PET, HiCAR, and related methods each yields a two-dimensional (2D) pairwise contact matrix encoding chromatin interaction frequencies and a one-dimensional (1D) coverage vector, indicating protein binding or chromatin accessibility. However, a lack of accessible and interpretable computational tools to quantify the similarity between these datasets prevents rigorous data quality assessment and biological implications.

**Results:** We developed HiChIA-Rep, an efficient algorithm incorporating both 1D and 2D signals to measure similarity via graph signal processing methods. Implemented in Python, HiChIA-Rep takes standard file format as inputs: .hic, .bedpe for 2D contact matrix and .bigwig or .bedgraph for 1D coverage vector. HiChIA-Rep utilizes 1D and 2D signals effectively to distinguish biological replicates from non-replicates, cell lines, experiment types, and protein factors, often outperforming existing tools. In addition, HiChIA-Rep can evaluate specific genomic regions of interest, and its utility is demonstrated for correlating differential gene expression and transcriptional chromatin interactions mediated by RNA Polymerase II in MCF10A and MCF7 cell lines.

**Conclusions:** HiChIA-Rep is readily applicable to existing ChIA-PET, HiChIP, PLAC-seq, HiCAR, and ChIATAC data. More broadly, HiChIA-Rep provides a graph signal processing framework to jointly analyze 1D and 2D signals, paving the ways to incorporate and interpret multiple data modalities probed by the emerging single-cell multi-ome methods in the future. We expect HiChIA-Rep to be a fundamental tool for the 3D genomics community.

## BACKGROUND

The three-dimensional (3D) genome organization is recognized as an important aspect of gene regulation and cellular functions (Bonev and Cavalli, 2016). To probe structures in genome folding, researchers developed several high-throughput sequencing technologies that rely on proximity ligation to link genomic loci that are physically proximal but far apart along the linear DNA sequence. Hi-C (Lieberman-Aiden et al., 2009; Rao et al., 2014), Micro-C (Hsieh et al., 2015) and related methods yield genome-wide unbiased contact frequencies (**Figure 1a**, left panel). In contrast, ‘enrichment-based’ technologies such as ChIA-PET (Fullwood et al., 2009; Wang et al., 2021), HiChIP (Mumbach et al., 2016), and PLAC-seq (Fang et al., 2016) employ a chromatin immunoprecipitation (ChIP) step to obtain chromatin interactions that are mediated by a specific protein factor of interest: CTCF, the CCCTC-binding factor (blue) (**Figure 1a**, middle panel), the cohesin subunit RAD21 (green), or RNAPII (RNA Polymerase II) (purple) to isolate contacts associated with transcriptional activity. More recently, methods utilizing Tn5 transposase—Trac-looping (Lai et al., 2018), OCEAN-C (Li et al., 2018), HiCAR (Wei et al., 2022), and ChIATAC (Chai et al., 2023)—were effective in capturing chromatin contacts among open chromatin regions (**Figure 1a**, right panel).

**Figure 1:**
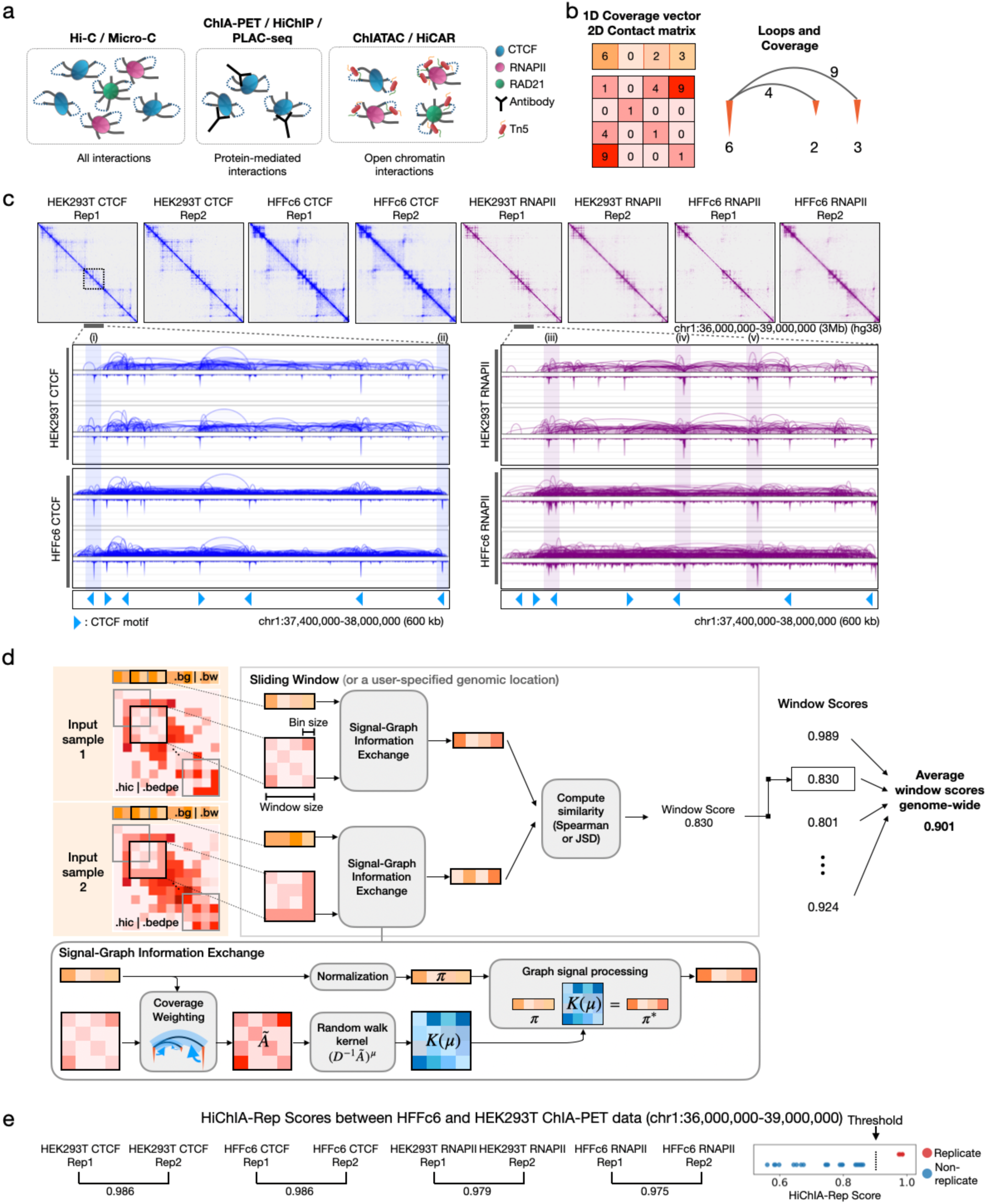
Overview of enrichment-based 3C technologies and HiChIA-Rep method. **(a)** Comparison of 3C assays. Hi-C and Micro-C capture digested and ligated DNA fragments which may be bound to various proteins. Enrichment-based technologies capture a subset of these products that either specifically interact with a protein factor (ChIA-PET, HiChIP and PLAC-seq) or interact with open chromatin regions (ChIATAC and HiCAR). CTCF: CCCTC-binding factor, RNAPII: RNA Polymerase II. **(b)** Data representations of enrichment-based chromatin interaction data. Left: 1D coverage vector and 2D contact matrix. Right: 2D contact matrix visualized as loops (gray) and coverage vector visualized as inverted peaks (orange) **(c)** Region chr1:36,000,000-39,000,000 of CTCF and RNAPII ChIA-PET data on HEK293T and HFFc6 human cell lines. Inset shows loops and coverage view and CTCF motifs track with light blue arrows indicating the binding motif orientation. Blue highlights (i) and (ii) show a peak with different binding signal strength between HEK293T CTCF and HFFc6 CTCF ChIA-PET data, while purple highlights (iii)-(v) show peak with different locations and differential loops between HEK293T RNAPII and HFFc6 RNAPII ChIA-PET data. Each dataset has two replicates. **(d)** HiChIA-Rep schematics. The program takes two input samples’ coverage vector (.bigWig or .bedGraph) and contact matrix (.hic or .bedpe) and computes the mean similarity of a sliding window across the binned genome. The ‘Signal-Graph Information Exchange Module’ (bottom) emphasizes loops with peak support and then performs a *μ*-step random-walk with the kernel *K*(*μ*) = (*D*^−1^*Ã*)*^μ^*, where *Ã* is the adjacency matrix and *D* is the diagonal matrix of its row sums (**Methods**). **(e)** Left: HiChIA-Rep window scores (chr1:36-39Mb) using default parameters (*μ* = 5, Spearman similarity) between replicates of HEK293T and HFFc6 ChIA-PET data. Right: Dot plot of HiChIA-Rep window scores (chr1:36-39Mb) between all pairwise combinations of HEK293T and HFFc6 ChIA-PET data. Each dot corresponds to a similarity score between either replicates (red) or non-replicates (blue). The threshold is drawn at 0.9.

Enrichment-based techniques yield multi-modal data: (1) a two-dimensional pairwise contact matrix where the *ij*-th entry corresponds to the number of interactions between binned genomic loci *i* and *j* as in Hi-C experiments (‘2D contact matrix’), and (2) one-dimensional coverage vector whose entry denotes the protein binding affinity or DNA accessibility at a particular location along the genome, as in ChIP-seq or ATAC-seq data (‘1D coverage vector’) (**Figure 1b**, left panel). An alternative representation of the data is the ‘loops and coverage’ view, where the ‘loops’ is defined by two genomic coordinates (‘loop anchors’) with a paired-end tag count (i.e., chromatin interaction frequency). The ‘loops’ is a file obtained from ChIA-PIPE (Lee et al., 2020) or other pipelines (e.g., ChIA-PET tool, Li et al., 2010; MAPS, Juric et al., 2019) and is not equivalent to the ‘chromatin loops’, which are dot-like patterns computationally identified from Hi-C contact maps. The ‘coverage’ refers to the 1D coverage vector shown as inverted peaks (orange) and the loops are drawn as arcs between the peaks (**Figure 1b**, right panel). A commonality among enrichment-based 3C assays is the two data modalities that are generated – the 2D contact matrix and 1D coverage vector.

These experimental technologies have revealed the human genome to be tightly organized into compartments and domains (Rao et al., 2014; Tang et al., 2015) and that gene regulation is aided by physical contacts between enhancers and promoters (Li et al., 2012; Weintraub et al., 2017; Wei et al., 2022). In addition to independent labs, the ENCODE (Luo et al., 2020) and 4D Nucleome consortia (Dekker et al., 2017; Reiff et al., 2022; Dekker et al., 2023) have generated more than 280 ChIA-PET and related IP-based datasets, and more than 90 HiCAR and other assays enriching for open chromatin regions. Computationally efficient and reliable methods are needed to interpret these large-scale datasets.

A critical preliminary task when analyzing data is to determine the reproducibility between two enrichment-based chromatin interaction datasets. For instance, a similarity score reveals whether replicates or patient samples can be pooled for downstream analyses. Beyond reproducibility assessment, downstream analysis itself is often aided by a specialized similarity measure, required for low-dimensional embeddings of samples or to quantify subtle biological discrepancies in regions of interest (e.g., encompassing oncogenes or genetic variants). The success of these tasks depends on designing a similarity measure that can resolve joint patterns in the data modalities –1D coverage vector and 2D contact matrix. In addition, a method that can distinguish the contribution of each data modality separately will be important in understanding the biological implications of the similarity or differences in datasets.

Unfortunately, due to the lack of computational methods, the current practice relies on qualitative visual inspections of the loops and coverage tracks, which can result in highly variable and inconsistent assessments. Alternatively, one may apply the tools that are designed to measure similarity between two Hi-C replicates. Representative software packages include GenomeDISCO (Ursu et al., 2018), HiCRep (Yang et al., 2017), HiC-Spector (Yan et al., 2017), and QuASAR (Sauria and Taylor, 2017). However, our results indicate that these tools are not applicable to ChIA-PET data as they often assign similar or higher score to non-replicates than to replicates. Crucially, these existing methods do not utilize the one-dimensional coverage vector inherent in enrichment-based 3C data that are indicative of protein binding or open chromatin sites. Two methods were proposed to overcome this limitation. HPRep (Rosen et al., 2021) extended HiCRep by modeling binding intensities, while AREDCI (Sun et al., 2024) proposed a self-similarity-based algorithm. However, their utilities were limited to a particular experimental type and file format, often requiring users to process the data from raw FASTQ file using specific pipelines tailored for the experimental protocol (e.g., restriction enzymes for HiChIP and PLAC-seq; linker sequence for ChIA-PET). Therefore, the 3D genomics field is still missing a sophisticated algorithm implemented in a readily accessible software tool to compare datasets from any enrichment-based 3C assays in standard data formats.

To fill in this gap, we developed HiChIA-Rep—high-resolution chromatin interaction analysis via reproducibility assessment—a Python package for quantifying the similarity between two ChIA-PET or related datasets. Our central contribution is leveraging graph signal processing techniques to holistically evaluate the data modalities. We achieve this by viewing the normalized 2D contact matrix as a weighted graph and the coverage vector as a signal on the nodes. Between two samples, we compare the dispersion of the coverage vector in this structured space, thereby jointly assessing the differences in both modalities. HiChIA-Rep has two unique practical features lacking in existing tools. The first feature is the ability to compare specific genomic regions of interest for a comparison between two cell types. The second feature is to quantify the individual contribution of 1D coverage and 2D contacts through ablation tests, determining if a region exhibits similar coverage profiles, contact matrices, both, or neither. One potential application is distinguishing between an RNAPII-mediated interaction that bridges a promoter and distal enhancer and independent RNAPII binding at the two regions without physical contact. These scenarios may have similar RNAPII coverage profiles but different contact matrices.

We tested HiChIA-Rep and other existing tools on ChIA-PET, HiChIP, PLAC-seq, ChIATAC, HiCAR data in human, Drosophila melanogaster, and mouse cells. In particular, HiChIA-Rep can: (1) clearly distinguish biological replicates from non-replicates, (2) quantify biological differences between ChIA-PET libraries of variable conditions (e.g., wild type vs. treated), (3) identify sub-regions of the genome that are different or similar, (4) compute the pairwise similarities of multiple cell lines to construct the chromatin interaction phylogeny that is representative of developmental trajectories.

## RESULTS

### Overview of HiChIA-Rep

When comparing two ChIA-PET experiments, the data may differ in either the 2D contact matrix, the 1D coverage vector, both, or none. For instance, HEK293T and HFFc6 CTCF ChIA-PET contact matrices in a 3 Mb region on chromosome 1 appear to be correlated (**Figure 1c**), but a closer look into a 600kb subset of the region (inset) shows differences in the CTCF protein binding signals (**Figure 1c**, i-ii). For RNAPII ChIA-PET data, however, HEK293T and HFFc6 exhibit not only differing binding signals, but also loop structures as shown by (iii)-(v). This noted property motivated us to develop a reproducibility measure that can assess the similarity between two ChIA-PET experiments by incorporating the inherent correlations in the 2D contact and 1D coverage. There are currently limited reproducibility methods for enrichment-based sequencing data, and simple applications of existing methods, such as Hi-C reproducibility metrics, would ignore the coverage signals. Perhaps a more subtle issue of using Hi-C methods involves the underlying assumption that reads are uniformly sampled across the genome, which is often implicitly assumed by downstream algorithms (e.g. Hi-C normalization methods). By contrast, ChIA-PET data are not expected to have uniform coverage as they enrich for specific subsets of interactions along the genome. As a result, an important step in ChIA-PET preprocessing is to emphasize interactions (loops) that are supported by strong coverage vector signal. One approach is ‘peak support filtering’ that removes loops if the loop anchors do not coincide with peaks (**Figure S1a**). The peak support filtered loops remove noise and emphasize relevant biological interactions, such as CTCF loops for CTCF ChIA-PET (top) and enhancer-promoter (E-P) interactions for RNAPII ChIA-PET (bottom). However, this method relies on accurate set of peaks to define regions of high enrichment signal, requiring users to run a peak-calling software and assuming only a binary output (a peak exists or not). Instead, we perform ‘coverage weighting’, which adds the enrichment signals coinciding at the anchors and multiplies the sum to the respective loop weight, thereby emphasizing well-supported loops in a continuous, non-binary manner (**Figure S1b**).

Incorporating coverage weighting, we developed HiChIA-Rep, which can quantify the similarity between two datasets from ChIA-PET or related enrichment-based assays genome-wide or at a specific genomic region (**Figure 1d**, see **Methods** for details). Specifically, HiChIA-Rep takes two input samples’ contact matrices (.hic or .bedpe) and coverage vectors (.bedGraph or .bigWig) binned at a specified bin size, and performs a sliding window across the genome. Each window is fed into the ‘Signal-Graph Information Exchange’ module that is designed to pass information between the matrix and vector representations. Specifically, we construct a normalized contact matrix *Ã* using coverage weighting (**Figure S1b**). Then, we simulate a *μ*-step random walk of the coverage vector on the normalized graph (**Figure 1d** ‘Graph Signal Processing’) and compare the resulting distributions after random walk using Spearman or the Jensen-Shannon Divergence (JSD). A weighted average of window scores is taken for each chromosome, and the resulting chromosomal scores are then averaged to obtain the genome-wide similarity measure between the two input samples. The effect of these operations on real data, MCF7 CTCF ChIA-PET rep1, is provided (**Figure S1c**). Notably, HiChIA-Rep does not rely on complex training steps and is designed to be a principled signal processing method. Algorithmic details are provided in the **Methods** section, and the software is publicly available as a Python package.

### HiChIA-Rep can distinguish replicates from non-replicates in HEK293T and HFFc6 ChIA-PET data

Applying HiChIA-Rep to HEK293T and HFFc6 ChIA-PET data with two replicates each for CTCF and RNAPII, we find consistently high scores (>0.9) between replicates for the 3 Mb region shown in **Figure 1c** (**Figure 1e**). We also compared all possible combinations of non-replicate pairs for the same region and show that all non-replicate scores can be separated (<0.9) from the replicate scores (**Figure 1e**, right). We then sought to establish whether the HiChIA-Rep reproducibility scores can be used to cluster the samples correctly in terms of the cell line and factor. We show a pairwise comparison matrix of genome-wide HiChIA-Rep scores with hierarchical clustering between HEK293T and HFFc6 ChIA-PET data (**Figure S1d**). Here, HiChIA-Rep correctly clustered based on the factor (RNAPII vs CTCF), followed by the cell line, which aligns with the notion that CTCF binding and loops are conserved across cell lines and species (Klenova et al., 1993; Phillips and Corces, 2009; Moon et al., 2005; Kim et al., 2007; Rudan et al, 2015). Notably, RNAPII is known to be less conserved than CTCF across cell lines (Tang et al., 2015; Li et al., 2012; Kim et al., 2026), which is reflected in the HiChIA-Rep scores: HEK293T vs HFFc6 RNAPII (0.619-0.637) vs. HEK293T vs HFFc6 CTCF (0.757-0.790).

As a comparison, we also tested AREDCI and HPRep, which were designed to assess the reproducibility of ChIA-PET and PLAC-seq data, respectively. AREDCI is a comprehensive pipeline with a web interface that requires .MICC files from the ChIA-PET2 processing pipeline (Li G et al., 2017). We modified the ChIA-PET2 pipeline to take input loops files to supply at an intermediary step to convert into .MICC files, amounting to 1 hour 40 minutes to preprocess 8 samples (**Methods**). HPRep is an end-to-end pipeline starting with a bam file using MAPS, and is designed to work for PLAC-seq data. Because ChIA-PET utilizes AB linker (Fullwood et al., 2009) or bridge linker sequences (Li X et al., 2017) and PLAC-seq relies on biotin pull-down, computational pipelines differ in identifying the ligated sequences: the former searches for linker sequences while the latter for restriction enzyme sites. Thus, a viable way for us to run HPRep was to interject the program using .bedpe file containing a list of interacting loop pairs with 7^th^ column denoting PET counts (see **Methods**). AREDCI clustered the replicates well with its scores between 0.912 to 0.94, and non-replicates between 0.809 and 0.866 (**Figure S1e**, left panel). We note that the separation between largest non-replicate score and smallest replicate score is 0.046 for AREDCI and 0.14 for HiChIA-Rep. HPRep failed to distinguish replicates from non-replicates: a non-replicate score between HFFc6 CTCF Rep1 and HEK293T CTCF Rep 1 at 0.518 was higher than the replicate score between HFFc6 RNAPII at 0.509 (**Figure S1e**, right panel).

We then compared our method against four existing Hi-C reproducibility methods, HiCRep, GenomeDISCO, HiC-Spector, and QuASAR (**Figure S1f**) and found their clusters to prioritize cell line over protein factor. More importantly, the four methods failed to distinguish replicates from non-replicates, with non-replicates having higher reproducibility scores than replicates (**Figure S1f**, **Figure 2a**). The HiChIA-Rep threshold for reproducibility is 0.9 and is computed from extensive tests on various cell-lines and technologies (**Methods**). The total run times on the HFFc6 and HEK293T ChIA-PET dataset (28 pairwise comparisons) are reported for Hi-C reproducibility methods on a HPC environment: QuASAR had the shortest run time (6.5s), followed by HiCRep (130s), HiChIA-Rep (45 minutes), GenomeDISCO (136 minutes), and HiC-Spector (159 minutes) (**Figure 2b**, left). Due to AREDCI having a graphical user interface, we used a local machine to measure run-times for enrichment-based data reproducibility methods: AREDCI (15s), HPRep (37 minutes), and HiChIA-Rep (41 minutes) (**Figure 2b**, right).

**Figure 2:**
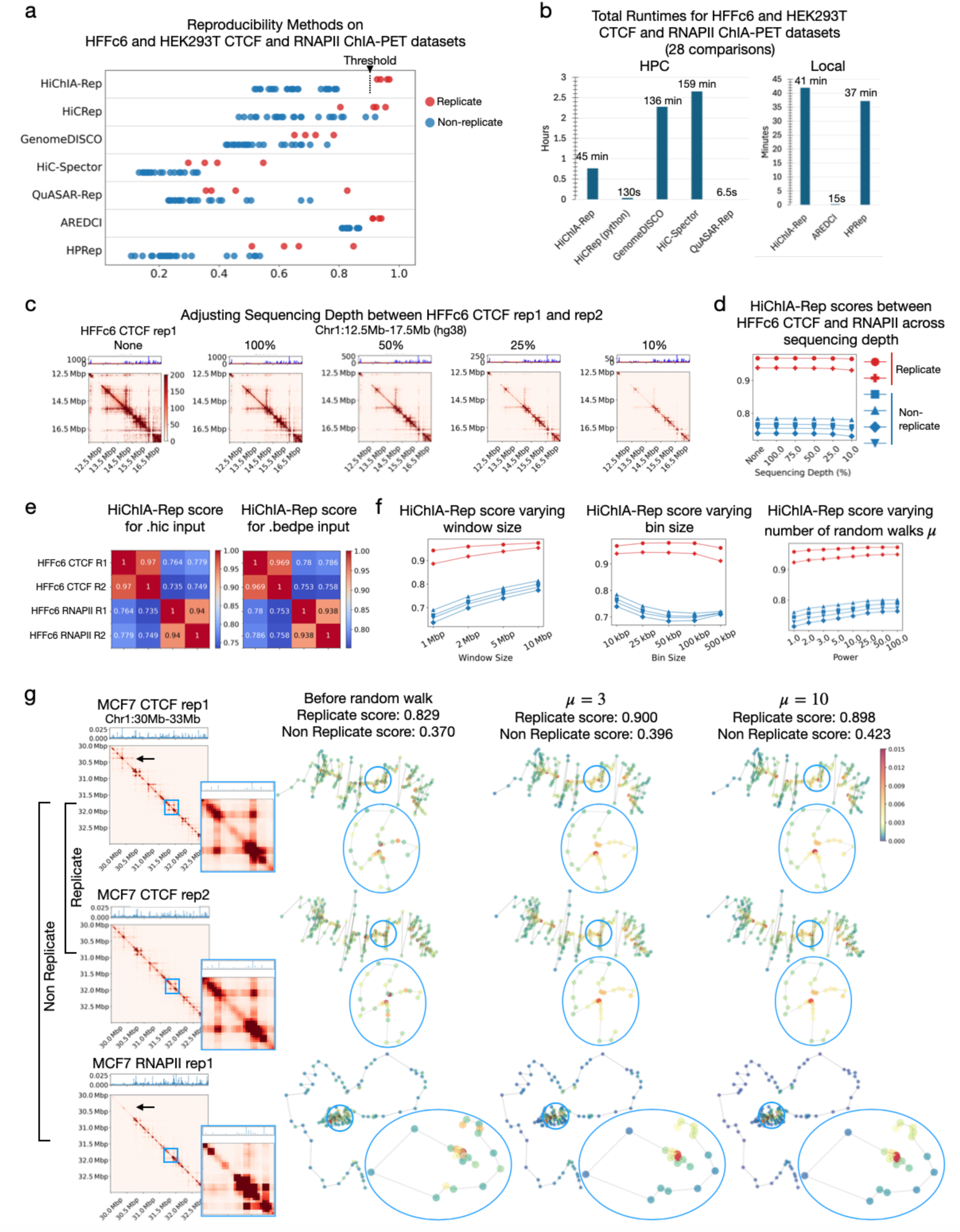
Comparison of HiChIA-Rep against existing reproducibility methods and across sequencing depth and input parameters. **(a)** Dot plot of HiChIA-Rep, Hi-C reproducibility, and AREDCI and HPRep scores between all pairwise combinations of HEK293T and HFFc6 ChIA-PET data. Threshold is drawn at 0.9 for HiChIA-Rep. **(b)** Total runtimes of HiChIA-Rep and other reproducibility methods on HEK293T and HFFc6 ChIA-PET data (encompassing a total of 6 pairs) in HPC environment using 4 core and 64GB RAM (Left) and local environment using 10 core CPU and 32GB RAM (Right). **(c)** 2D contact matrices and 1D coverage vector of HFFc6 CTCF rep1 (replicate 1) adjusted for sequencing depth with HFFc6 CTCF rep2. Sequencing depth adjustment: no sequencing depth adjustment (None); downsampled such that total number of counts is equal to the smaller total count in a given pair (100%), and further downsampling (50%, 25% and 10%). **(d)** HiChIA-Rep scores on HFFc6 CTCF and RNAPII ChIA-PET data varying sequencing depth, with replicate scores in red and non-replicate scores in blue. **(e)** HiChIA-Rep scores on HFFc6 CTCF and RNAPII ChIA-PET data for .hic input (left) and .bedpe input (right). **(f)** HiChIA-Rep scores on HFFc6 CTCF and RNAPII ChIA-PET data varying window size (left), bin size (center), and number of steps in random walk (right) **(g)** Left: Coverage vector and contact matrix of MCF7 CTCF (rep1, rep2) and RNAPII rep1 ChIA-PET data on chr1:30-33Mb region. Inset (blue border) shows chr1:31,750,000-32,050,000. Right: MDS embedding of contact matrix with color of points showing coverage vector before random walk, after random walk (*μ* = 3), and after random walk (*μ* = 10). The scores are the Spearman coefficient for the chr1:30-33Mb window.

### HiChIA-Rep is robust to sequencing depth and parameters

To gauge the effect of sequencing depth on HiChIA-Rep, we used a binomial downsampling scheme to ensure that the number of total counts between two input samples is matched to the lower count of the pair for both the contact map and the enrichment signal (**Figure 2c**). We generated sub-sampled pairs for HFFc6 CTCF (rep1 and rep2) and RNAPII (rep1 and rep2) ChIA-PET data. Applying HiChIA-Rep on this dataset resulted in scores that remain consistent across sequencing depth ranging from 100 percent to 10 percent (**Figure 2d**). We then tested the effect of various input types and parameter settings. HiChIA-Rep can take as input either a .hic or .bedpe file for the pairwise chromatin contacts, between which results are largely consistent (**Figure 2e**). This feature is critical to ensure that HiChIA-Rep is a general and versatile tool for users with any standard file format without having to re-process datasets from raw reads. We then tested the three main parameters of HiChIA-Rep: window size, bin size, and the number of steps *μ* in random walk. We varied window size from 1 Mb to 10 Mb, bin size from 10 kb to 500 kb, and *μ* from 1 to 100. Unless otherwise specified, the other parameters were kept as default parameters specified in the **Methods** section. The separability of replicates and non-replicates remained stable across the set of parameters (**Figure 2f**). Interestingly, there appears to be an optimal bin size at around 50 kb or 100 kb (**Figure 2f**, center), suggesting a tradeoff between noise at small bin sizes and a lack of information at large bin sizes. We see an increase in run-time with window size 10 Mb (29 minutes compared to about 22 minutes for the default of 5 Mb), whereas other parameter variation did not significantly affect run-time (**Figure S2a**). We also evaluated whether HiChIA-Rep scores using JSD as the comparison metric was reliable across input types (**Figure S2b**) and parameters (**Figure S2c**). All results across input types and varying parameters ran with peak memory consumption under 30 GB.

To investigate the effect of increasing *μ* on HiChIA-Rep scores, we show a replicate and a non-replicate pair of MCF7 ChIA-PET data in a 3 Mb window on chromosome 1 (**Figure 2g**). We selected this region as there were major structural differences between CTCF and RNAPII data, as shown in the inset. We embedded the contact matrix graph in 2D using multidimensional scaling (MDS), and colored each node according to the coverage vector. Focusing on a chromatin loop (inset) that is present in the CTCF data (rows 1 and 2) but not in the RNAPII data (row 3), we find that the random walk operation (*μ* = 3 and *μ* = 10) smooths the coverage vector to nodes adjacent along the linear genome. To further isolate the effect of the random walk on the coverage vector, we ablated the vector to be a delta-like spike for both CTCF and RNAPIl (**Figure S2d**). Since the signals are identical, the similarity scores before random walk are 1. With random walk (*μ* = 3 and *μ* = 10), however, we find that the signal disperses according to the topology of the chromatin graph: for CTCF at *μ* = 10, the delta spike originally placed on genomic locus A has spread through a chromatin loop and signal has accumulated on locus B. Comparatively, RNAPII does not have this chromatin loop and the signal has not dispersed in the same pattern (**Figure S2d**, row 3). As *μ* increases, the coverage vector gains structural information of nodes that are close in 3D nuclear space (e.g. between the chromatin loop). Indeed, this behavior can be analyzed within the framework of Markov Chains, where the limiting distribution will depend only on the normalized contact matrix and not the initial coverage vector under regularity conditions.

### HiChIA-Rep can compare ChIA-PET, HiCAR, ChIATAC, and HiChIP data across 8 cell lines

We next applied our method on an extended ChIA-PET dataset consisting of four human cell lines HFFc6, MCF7, MCF10A, and HEK293T enriched for CTCF and RNAPII with two replicates each (**Figure 3a**). HiChIA-Rep can distinguish protein factors, cell line and replicates from non-replicates (**Figure 3a**). Scores are generally higher for CTCF than for RNAPII (**Figure S3a**), reflecting the conserved nature of CTCF binding and looping across cell lines, whereas RNAPII-associated transcriptional patterns suggest a role that is specific to a cell line. We noticed that HFFc6 CTCF has high correlation with HFFc6 RNAPII (0.735-0.779) as compared to other CTCF-RNAPII similarity scores (0.402-0.660) (**Figure 3a**), potentially due to the presence of spurious interactions present in the loops. Nevertheless, HiChIA-Rep can separate all replicates from non-replicates (**Figure 3a**).

**Figure 3:**
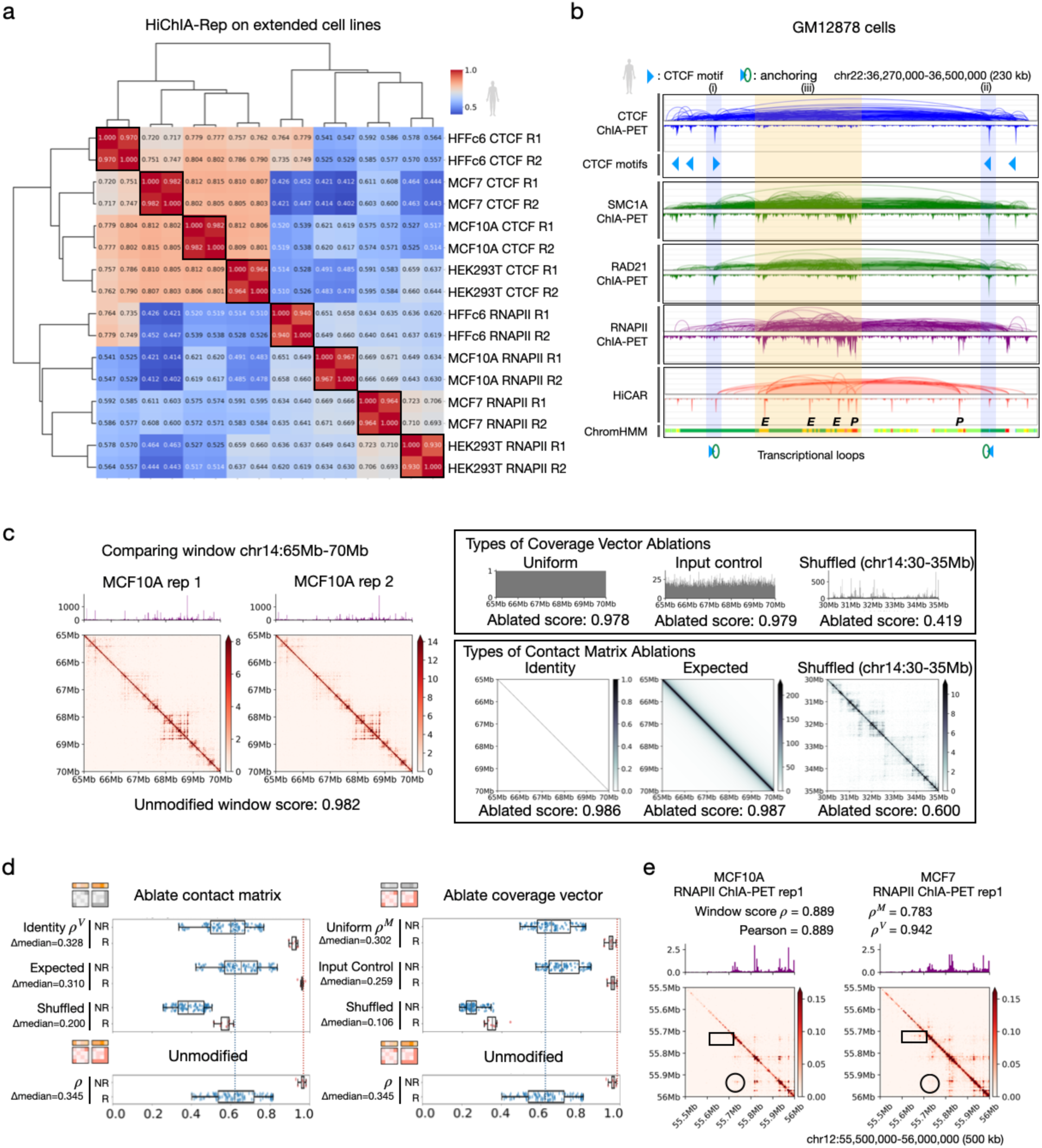
HiChIA-Rep on human ChIA-PET data and GM12878 HiCAR. **(a)** HiChIA-Rep scores between all pairwise combinations of HFFc6, MCF7, MCF10A, HEK293T cell lines for CTCF and RNAPII ChIA-PET (2 replicates each) with hierarchical clustering. ‘R’ denotes replicate. **(b)** Genome browser tracks showing GM12878 data on chr22:36,270,000-36,500,000: CTCF ChIA-PET ‘loops and coverage’ (row 1), CTCF motifs (row 2), SMC1A, RAD21, and RNAPII ChIA-PET (rows 3-5) and HiCAR (row 6), and ChromHMM markings (row 7). Active promoters (‘P’; red), gene transcription (dark green), enhancers (‘E’; yellow). Blue highlight (i) and (ii) shows the anchors of a cohesin-CTCF loop. Orange highlight (iii) shows densely clustered enhancer and promoters. **(c)** Left: An example window showing coverage vector and contact matrix of chr14:65Mb-70Mb MCF10A RNAPII ChIA-PET rep1 and rep2 data. Right: Different ablation types for coverage vector (uniform, input control, shuffled) and contact matrix (identity, expected, shuffled) (**Methods**). The ‘ablated score’ is the window score after replacing the coverage vector or contact matrix with the respective ablation. **(d)** HiChIA-Rep scores on different ablation experiments for contact matrix (left) and coverage vector (right) on extended cell lines dataset (HFFc6, MCF7, MCF10A, HEK293T cell lines for CTCF and RNAPII ChIA-PET with 2 replicates each). Last row is the unmodified program. ‘*Δ*median’ is the difference of median of replicate scores ‘R’ and non-replicate scores ‘NR’. **(e)** An example region on chr12 showing window scores comparing MCF10A and MCF7 RNAPII ChIA-PET: unmodified *ρ*, window score of contact matrix only *ρ^M^*, window score of coverage vector only *ρ^V^* and Pearson correlation coefficient between coverage vectors.

Next, we looked at GM12878 cell line as it has a large profile of genomics data. Here we show ChIA-PET data for CTCF, RNAPII, SMC1A, and RAD21, where SMC1A and RAD21 are both subunits of cohesin and thus considered as replicates (**Figure 3b**). We also show HiCAR data that captures open chromatin associated contacts. Highlights (i) and (ii) show two CTCF peaks with convergent motifs and an overarching loop between them, suggesting their role as anchors of a chromatin loop. The bottom two tracks show RNAPII and HiCAR data, which are correlated in region (iii) containing enhancer and promoters as defined by ChromHMM (Ernst and Kellis, 2012), consistent with the fact that open regions tend to be transcriptionally active (Bell et al., 2011). Interestingly, the enhancer-promoter loops are moderately present in the cohesin tracks (RAD21 and SMC1A) but absent in the CTCF, further supporting an emerging notion that cohesin may play a role in some of these enhancer-promoter interactions whereas CTCF is strictly associated with the cohesin-CTCF chromatin looping mechanism (Kim et al., 2026; Guckelberger et al., 2026; Kane et al., 2022). As the authors of HiCAR have noted (Wei et al., 2022), some CTCF binding and looping is captured by the HiCAR data, as shown by the blue highlighted region.

We wanted to test whether the aforementioned relationships among protein factors and open regions could be captured using HiChIA-Rep. Accordingly, HiChIA-Rep has highest correlation between the cohesin tracks SMC1A and RAD21 (0.947) (**Figure S3b**), which is followed by correlation between cohesin tracks and CTCF (0.893, 0.925). We find that RNAPII is more similar to cohesin (0.854, 0.823) than CTCF (0.725) as expected. While HiCAR is more distant to ChIA-PET datasets, HiCAR data appears to be equally similar to RNAPII (0.711) and cohesin (0.708, 0.716), which is plausible as the E-P interactions and cohesin looping are both captured by HiCAR (**Figure 3b**). HiChIA-Rep captured the patterns that RNAPII and CTCF ChIA-PET are the most distant among ChIA-PET datasets, while cohesin exhibits loops and bindings that resemble both CTCF (forming structural loops) and RNAPII (mediating transcriptional loops).

We then sought out whether HiChIA-Rep could yield consistent results across cell lines and other enrichment-based technologies. Here, we find separability of replicates and non-replicates on HiCAR and ChIA-PET data for GM12878 and H1 cell lines (**Figure S3c**). We then applied HiChIA-Rep on *Drosophila melanogaster* S2 ChIATAC data, which is a similar enrichment-based method to HiCAR in selecting chromatin contacts associated with open regions (Chai et al., 2023) (**Figure S3d**). HiChIA-Rep failed to distinguish replicates between RNAPII ChIA-PET, possibly reflecting the poor ChIA-PET data quality as also reported by the authors of ChIATAC (Chai et al., 2023). On human HG00731 HiChIP data, HiChIA-Rep can distinguish replicates from non-replicates (**Figure S3e**). We then used HiChIA-Rep window scores to select two regions that had low similarity (left) and high similarity between HiChIP CTCF and SMC1 (**Figure S3f**). With the low similarity region (chr13:85Mb-90Mb) with a score of 0.653, we observed clear structural differences, such as a stripe and additional binding signal in the SMC1 data that is not present in the CTCF. By contrast, in the high similarity region (chr14:60Mb-65Mb) with a score of 0.93, both the structure and binding affinity are highly correlated between CTCF and SMC1. This example suggests that HiChIA-Rep scores depend on both the 1D coverage and 2D contacts, and the extent to which each contributes to the score is investigated next.

### Both 1D coverage and 2D contacts are necessary for accurate similarity measures

To investigate the quantitative contribution of each modality to the observed HiChIA-Rep performance, we performed ablation tests. Specifically, a 5 Mb region on chromosome 14 of MCF10A RNAPII ChIA-PET rep1 and rep2 shows a high degree of similarity between both the coverage vector and contact matrix, evidenced by high HiChIA-Rep score of 0.982 (**Figure 3c**, left panel). Ablation testings can probe whether this high score is a result of similar coverage vector or contact matrix. We considered 3 types of ablations for the coverage vector and contact matrix: (1) uniform and identity, (2) input control and expected, and (3) shuffled, where either the coverage vector or contact matrix is shuffled to a random genomic location (**Figure 3c**, right panel) (see **Methods** for details). We observed that the first two types of ablations (uniform/identity and input control/expected) have a high similarity score despite the ablation of one modality: 0.978–0.987. The shuffled ablation shows the most drastic drop in performance at 0.419 and 0.600 for coverage vector and contact matrix, respectively. We next tested on not only replicates but non-replicate pairs to evaluate the performance under each ablation.

We performed ablation tests genome-wide for all pairwise combinations on the extended cell-lines ChIA-PET dataset (HFFc6, MCF7, MCF10A, HEK293T) (**Figure 3a**). We observed that ablating the contact matrix results in lower separation between replicates and non-replicates than the unmodified program (incorporating both contact matrix and coverage vector) (**Figure 3d**, left panel). For example, the difference in medians of replicate and non-replicate scores was 0.345 in the unmodified, which dropped to 0.328, 0.31, and 0.2 for each of identity, expected, and shuffled models of ablating contact matrix. Similarly, ablating coverage vector reduced the median differences to 0.302, 0.259, and 0.106 (**Figure 3d**, right panel). These results highlight that both contact matrix and coverage vector strengthen the ability in separating replicates from non-replicates. The strict decrease in performance with all performed ablations implies that both components are necessary for the HiChIA-Rep program.

We used ablation tests to not only determine the necessity of the coverage vector and contact matrix, but as another similarity measure. The contact matrix identity ablation can equivalently be viewed as a similarity score utilizing only the coverage vector (*ρ^V^*). Likewise, the uniform coverage vector ablation is interpreted as a similarity score comparing the contact matrix only (*ρ^M^*). Using the three scores (*ρ^V^*, *ρ^M^*, and unmodified *ρ*), we identified a 500 kb region on MCF10A and MCF7 (normal vs. breast cancer cell lines) that exhibit similar binding coverage profiles with a high coverage vector similarity *ρ^V^* of 0.942, supported by Pearson’s correlation coefficient of 0.889, but with differential contact matrix: a chromatin loop present only in the MCF10A data (circle) and a stripe in only MCF7 (rectangle) (**Figure 3e**). Accordingly, the contact matrix similarity *ρ^M^* is lower at 0.783 (**Figure 3e**). Similarly, another 500kb region on chromosome 9 shows high coverage similarity with high Pearson’s correlation coefficient of 0.967 and *ρ^V^* of 0.926, but low contact matrix similarity *ρ^M^* at 0.791 (**Figure S3g**). Inspecting the contact matrices reveals a series of loops diminished in MCF7 (black circles) (**Figure S3g**, bottom left). These two examples illustrate that two regions may have high similarity in binding profile yet display notable structural differences in the contact matrices.

### HiChIA-Rep scores correlate with differential genes between MCF10A and MCF7

We next sought to further explore the discrepancies between MCF10A and MCF7 RNAPII ChIA-PET data (**Figure 3e, Figure S3g**) and whether the HiChIA-Rep scores correlate with transcriptional activity. To do so, we used public RNA-seq data with three replicates each of MCF10A and MCF7 (Barutcu et al., 2015) from which DESeq2 identified 1847 downregulated, 1932 upregulated, and 7857 unchanged genes (**Figure 4a**). Changes in gene expression, as measured by the mean of absolute log2 fold-change of genes in a window, were negatively correlated with HiChIA-Rep scores across the genome (Spearman’s correlation coefficient of - 0.209), suggesting that windows with minimal gene expression changes were likely to have similar RNAPII contacts and binding (**Figure 4b**, blue dots). As a control, MCF10A replicates had minimal changes in gene expression and high HiChIA-Rep scores (**Figure 4b**, red dots). DESeq2 gene classes also suggest downregulated and upregulated to be less similar than the unchanged class (**Figure 4c**). There was a statistically significant difference in gene expression between window scores 0.73–0.95 and 0.63–0.73 (p=0.0064), where lower differential gene expression is correlated with high HiChIA-Rep similarity between MCF10A and MCF7 (**Figure S4a**). Notably, transcriptional patterns remain constant across the three groups when comparing MCF10A replicates (**Figure S4b**).

**Figure 4:**
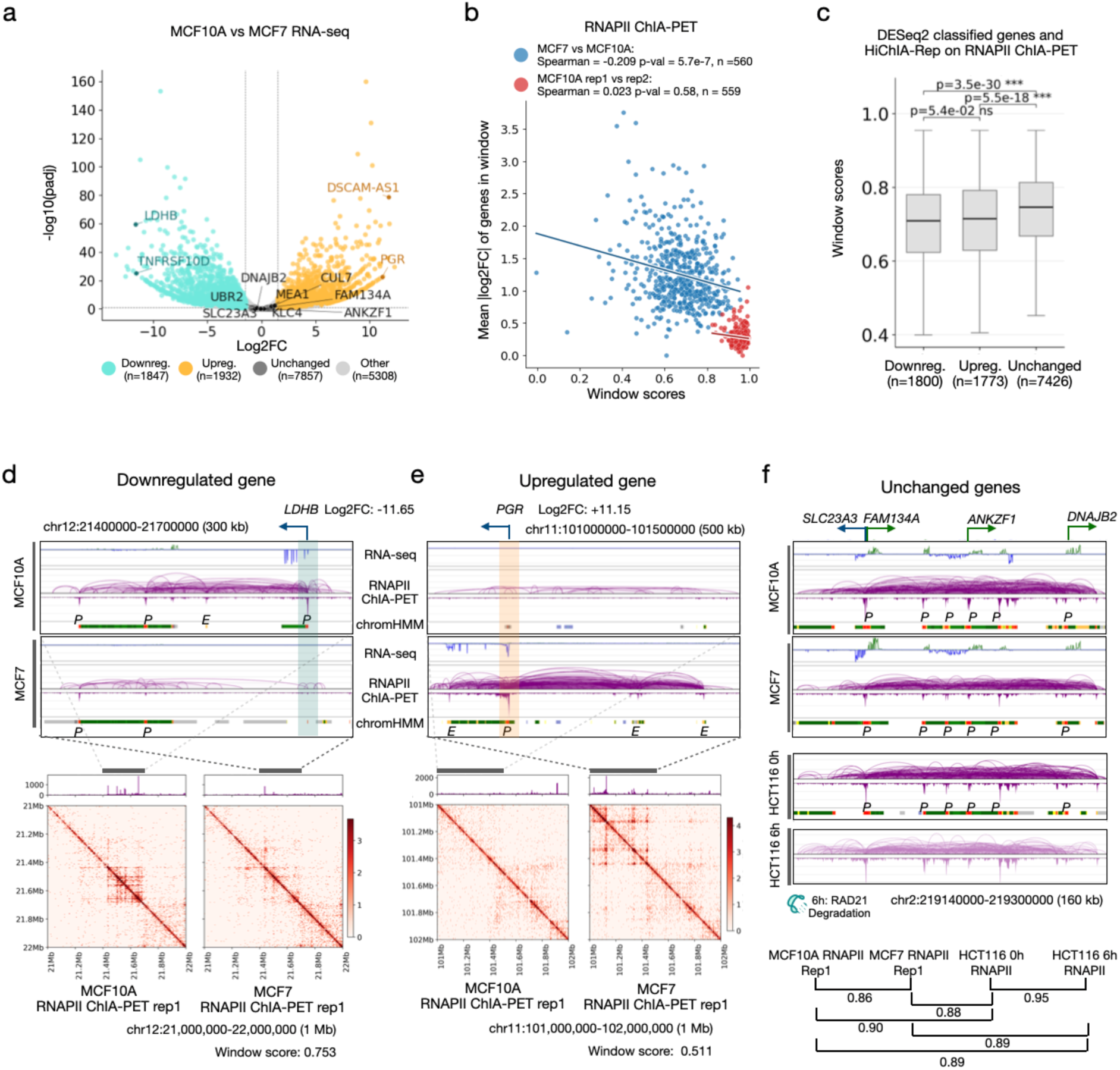
Correlating differential gene expression with HiChIA-Rep scores of RNAPII ChIA-PET data in MCF10A and MCF7 cells. **(a)** Volcano plot of differential genes between MCF10A and MCF7 using DESeq2 results on RNA-seq data. Each gene is classified into ‘downregulated’ (cyan), ‘upregulated’ (orange), ‘unchanged’ (dark gray), and ‘other’ (gray) genes. **(b)** Scatter plot of mean absolute log2 fold-change (FC) of genes in window and HiChIA-Rep window score comparing RNAPII ChIA-PET data: MCF7 and MCF10A (blue points) and MCF10A rep1 vs rep2 (red points). P-values are from two-sided Spearman’s rank correlation test. **(c)** Boxplot of genes classified using DESeq2 into ‘downregulated’, ‘upregulated’, and ‘unchanged’ and corresponding HiChIA-Rep window score comparing MCF7 and MCF10A RNAPII ChIA-PET. P-values are from two-sided Mann-Whitney U test. **(d)** Example of a downregulated gene *LDHB* in MCF7 with RNA-seq, ChIA-PET ‘loops and coverage’, and chromHMM states (‘P’ promoter in red, ‘E’ enhancer in yellow) for MCF10A and MCF7 (top). Bottom: Coverage vector and contact matrix of a larger genomic range encompassing the *LDHB* gene with HiChIA-Rep window similarity score. **(e)** Similar to panel **(d)** but showing an upregulated gene *PGR*. **(f)** Top: Browser visualization of select unchanged genes on chr2, showing RNA-seq, ChIA-PET ‘loops and coverage view’ and chromHMM states for MCF10A and MCF7. Middle: Same region showing ChIA-PET and chromHMM states for HCT116 0hr (no auxin) and 6hr (auxin treatment for RAD21 depletion) data. Bottom: HiChIA-Rep window scores of same region between all pairwise combinations of MCF10A, MCF7, HCT116 0hr, and HCT116 6hr RNAPII ChIA-PET data.

Accompanying the genome-wide patterns, we focus on an individual examples of downregulated, upregulated, and unchanged genes. One of the downregulated genes, *LDHB* (**Figure 4a**) (log2 fold-change of -11.65) has its promoter connected to a distal enhancer and other active promoters in MCF10A; by contrast, the enhancer mark is replaced by heterochromatin state in MCF7 and the binding and looping structures around *LDHB* diminished, quantified by the low HiChIA-Rep score of 0.753 (**Figure 4d**). A similar example is shown for *TNFRSF10D* with HiChIA-Rep score 0.706 (**Figure S4c**). Upregulated gene *PGR* (**Figure 4a**) with a log2 fold-change of 11.15, displays the opposite pattern: reduced binding and looping in MCF10A, and strong promoter mark connected to distal enhancers upstream and downstream in MCF7, resulting in a similarly low HiChIA-Rep score of 0.511 (**Figure 4e**). Another gene *DSCAM-AS1* (**Figure 4a**) has binding signal and looping in MCF7 but no RNAPII signatures in MCF10A; this region has the lowest HiChIA-Rep score out of the examples at 0.15 (**Figure S4d**). Finally, unchanged genes were accompanied by minimal changes in RNAPII binding and looping with a relatively high HiChIA-Rep score of 0.86 (**Figure 4f**). Interesting, these genes were connected to short-range loops (<50kb), similar to the unchanged genes in HCT116 cells before (0h) and after (6h) depleting a cohesin subunit RAD21 via auxin-inducible degron system (Kim et al., 2026). Indeed, the RNAPII ChIA-PET data in HCT116 cells 0h and 6h of auxin treatment in the same 160 kb region visually displays similar levels of RNAPII loops and binding, supported by high HiChIA-Rep scores between the four datasets ranging from 0.86 to 0.95 (**Figure 4f**). Another set of unchanged genes are also connected via promoter-centric loops and binding that are invariant across MCF10A, MCF7, HCT116 0h and 6h RNAPII ChIA-PET data (**Figure S4e**).

These genome-wide statistics and individual examples support the notion that changes in gene expression between two samples are correlated with the changes in RNAPII binding and looping structures, which we quantify through HiChIA-Rep. While HiChIA-Rep does not offer statistical confidence to the differential contacts and binding, it is still capable of measuring the overall similarity in a given region, which is a unique feature that is lacking in many reproducibility methods. We next investigate the relationship between genome folding and gene transcription in mouse development through time-series data with multiple tissue samples.

### HiChIA-Rep captures biological patterns in PLAC-seq data on developing mouse embryo tissues

Another variation of ChIA-PET and HiChIP is PLAC-seq. Enriched for H3K4me3-associated chromatin contacts, publicly available PLAC-seq data (Yu et al., 2025) were generated on developing mouse embryo tissues in time-series from embryonic day 12.5 (E12.5) to postnatal day 0 (P0). We first applied our method by fixing the timepoint to E12.5 to test whether HiChIA-Rep could capture tissue similarities. We find that the brain tissues (midbrain, hindbrain and forebrain) had the highest similarity amongst each other (**Figure 5a**), whereas limb and liver were most distant to the other tissues. We then looked at a 2 Mb region on chromosome 2 (chr2:70,000,000-72,000,000) for forebrain (FB), craniofacial prominence (CF), limb (LM), and liver (LV) at E12.5 along with RNA-seq data in the corresponding tissues (**Figure 5b**). While the gene expression profile in this region is relatively similar between tissues, we observed large structural differences in the loop strengths and binding intensities of H3K4me3 PLAC-seq data with high correlation between CF and FB. This pattern can also be seen in the contact matrix view (**Figure 5c**), where CF-FB have similar structures; LM-LV appear correlated, sharing stripes (i) and (ii). The HiChIA-Rep window scores reflect these correlations (**Figure 5d**), where the CF-FB pair is the highest (0.919) followed by LM-LV (0.833). Interestingly, HiChIA-Rep ranks the FB-LV pair as the least similar out of all comparisons at 0.651. This finding was surprising considering that the 1D coverage signals appear correlated (**Figure 5b**). Nevertheless, the RNA-seq track shows significant differences (i)-(iv) between LV and FB, more so than for the other pairs, suggesting that such a ranking is consistent with gene transcription for this region. Taken together, HiChIA-Rep can measure discrepancies without solely relying on the coverage vector or the contact matrix, providing a relative ordering that is consistent with local structures.

**Figure 5:**
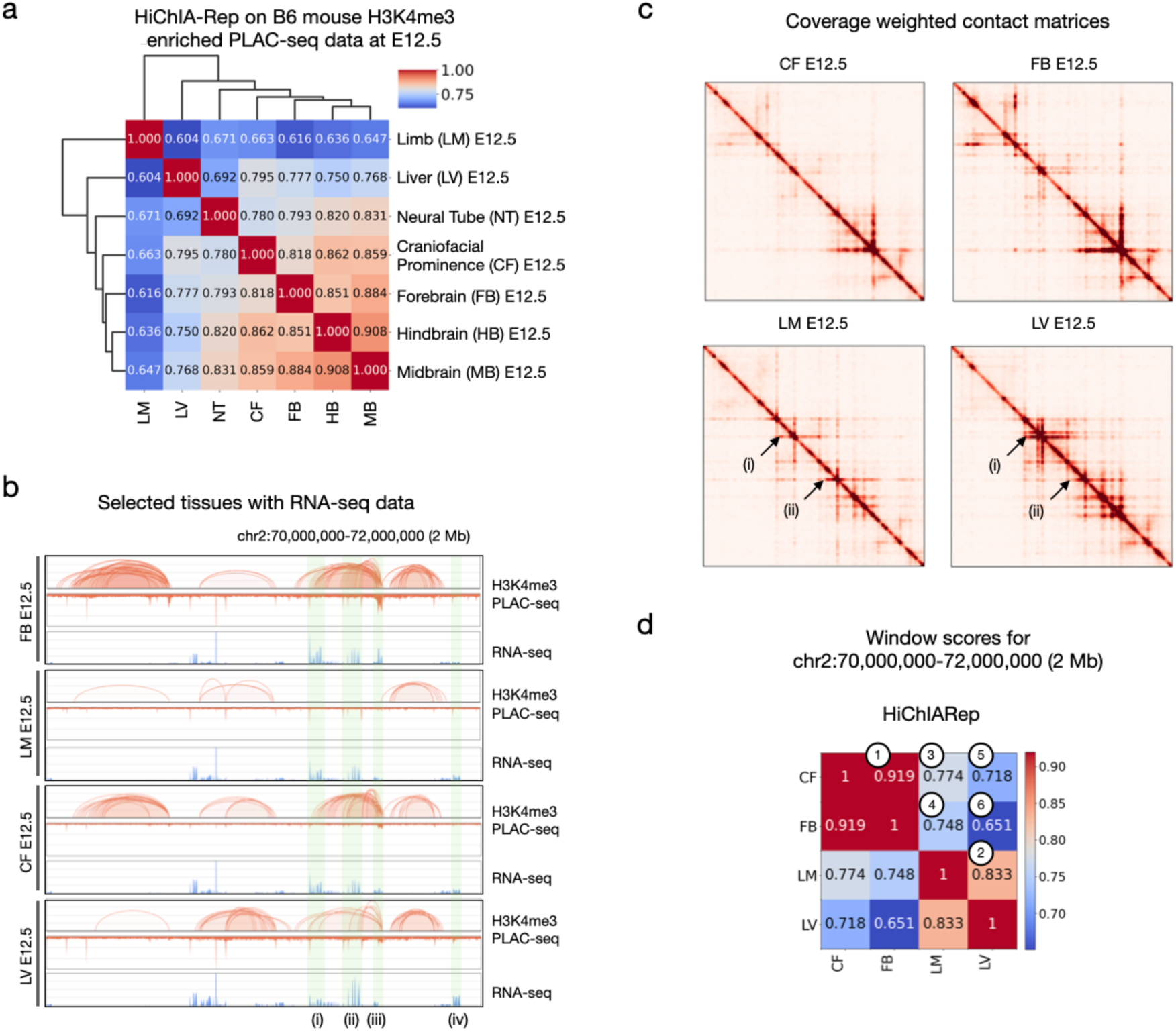
HiChIA-Rep on PLAC-seq data in developing mouse embryo tissue. **(a)** Hierarchical clustering of HiChIA-Rep scores when applied to PLAC-seq data enriched for H3K4me3 on B6 mouse embryo tissues at E12.5 (embryonic day 12.5). **(b)** Browser tracks showing mouse embryo tissues on chr2:70,000,000-72,000,000: PLAC-seq ‘loops and coverage’ and RNA-seq for forebrain (FB) (row 1), limb (LM) (row 2), craniofacial prominence (CF) (row 3), and liver (LV) (row 4). The green regions (i)-(iv) highlight differences in RNA-seq between the FB E12.5 and LV E12.5 pair. **(c)** Coverage weighted contact matrices at 10 kb resolution of the same samples shown in panel **(b)**. The arrows (i) and (ii) show shared stripes between limb (LM) E12.5 and liver (LV) E12.5. **(d)** HiChIA-Rep window scores on chr2:70,000,000-72,000,000. The number in the circles denotes the ranking from largest to smallest scores.

We then analyzed whether HiChIA-Rep could resolve the similarities between time-points (E12.5-P0) for the forebrain tissue, focusing on the *Neurod6* gene (chr6:55,400,000-56,000,000) involved in the differentiation of the nervous system (**Figure S5a**). Here, we see a gradual increase in transcription of *Neurod6* across the embryonic days and a slight reduction in the P0 sample. A similar trend is seen with PLAC-seq data, in which structures—loops and binding signals—appear and disappear across time (**Figure S5a**). This phenomenon is reflected in the HiChIA-Rep scores, where the P0 forebrain sample has a higher similarity to E13.5 than the other time-points (**Figure S5b**). When we used genome-wide HiChIA-Rep scores and embedded the similarity matrix in 2D coordinates, we find a similar pattern where P0 forebrain tissues cluster close to E13.5 or E15.5, suggesting that the dynamic program of epigenetic modification may not strictly align with time-points (**Figure S5c**). Applying HiChIA-Rep on the entire PLAC-seq dataset, we find that the major differences arise between the tissues more so than the time-points (**Figure S5d**), which was also apparent in the H3K4me3 ChIP-seq data presented in Yu et al., 2025. The HiChIA-Rep scores of PLAC-seq data also appears to be variable for certain samples (e.g., NT E12.5 rep1, LM E12.5 rep1) (**Figure S5d**), suggesting that those samples are worth additional investigation for quality assessment.

## DISCUSSION

We described HiChIA-Rep and demonstrated its utility in accurately quantifying the similarity between two datasets generated by a wide range of enrichment-based 3C assays. When tested on ChIA-PET, HiChIP, PLAC-seq, ChIATAC, HiCAR datasets in human, Drosophila melanogaster, and mouse cells, HiChIA-Rep distinguished replicates from non-replicates with high accuracy.

Our long-term goal is to understand the relationship between the 1-dimensional coverage vector (protein binding affinity or DNA accessibility) and 2-dimensional contact matrices (genome structure) and offer ways to incorporate information from both modalities. During the process, we will also extract the biological implications of these experimental protocols. For instance, the 1D coverage may be less prone to noise than 2D contacts for various reasons, such as the increased dimensionality of 2D matrices and potential random ligation noise. Additionally, we expect the 2D contact data to be inherently sparser than the enrichment signal, assuming that protein binding is required for loop formation: cohesin must be loaded onto the DNA in order for loop extrusion to occur, and similarly CTCF should be bound at the convergent motifs to stabilize this process. In other cases, the protein binding affinity may be noisy and uncorrelated with the 2D contacts when cells are not fixed properly during crosslinking, or when some protein factors move faster and spend less time on the DNA than others (e.g., RNAPII may exhibit higher mobility than CTCF, resulting in noisier-looking coverage tracks). We made our initial attempts in this direction through ablation tests in this study.

In terms of testing HiChIA-Rep on simulated data, there are some limitations in the current work. First, the method to adjust for sequencing depth is through a simple binomial distribution and may not be accurate as it is done after coverage weighting. An ideal approach would realistically model the sparsity of both 1D coverage and 2D contacts as if the same library were to be sequenced at a lower depth. Second, we did not test the robustness of HiChIA-Rep to noise, such as random ligation, non-specific pull-down during immunoprecipitation, and inefficient transposase activities. In both sequencing depth adjustment and noise simulations, we should also consider the distance-dependency of genome structure, where nearby DNA fragments are more likely to interact by chance than those far apart (Grob and Cavalli, 2017).

Finally, the distinction between local and global patterns becomes an important issue in analyzing PLAC-seq data generated from embryo tissues (**Figure 5**). Certain regions may exhibit a distinct tissue cluster configuration, driven by the activity of transcriptional elements and epigenetic modifications (**Figure S5a**), yet may not be representative of the “average” chromatin state across the genome (**Figure S5d**). As a result, local patterns that are meaningful but under-represented may be averaged out and therefore not captured by the (genome-wide) HiChIA-Rep score. Despite this limitation, the HiChIA-Rep outputs scores for individual windows of a sliding window, or a user-specified genomic region, which we used to our advantage in identifying regions of high or low similarity that are associated with pre-defined patterns (**Figure 3, 4**). A better approach to find and quantify the local similarities or discrepancies between samples is to identify statistically differential chromatin contacts with confidence measures, considering 1D signal, 2D contacts, and variability between replicates. We intend to pursue this line of work in the future.

## CONCLUSIONS

We present HiChIA-Rep, which has significant advances over existing approaches. Algorithmically, it applies a novel graph signal processing method to incorporate both 1D signals and 2D contacts present in enrichment-based 3C assays. Unlike other tools (Bhattacharyya et al., 2019; Lareau et al., 2018; Rosen et al., 2021), HiChIA-Rep does not require a set of peaks called from 1D signals and instead fully utilizes the tracks as continuous signal without arbitrarily setting a threshold to binarize signal as peaks or non-peaks. Being robust to various parameters and its algorithmic simplicity is a noted strength. Also, because the kernel operations rely on matrix-vector multiplication, the HiChIA-Rep software package is highly efficient in terms of runtime and may be further improved with the GPU acceleration in the future. Practically, HiChIA-Rep is a versatile, general, and user-friendly Python package. It takes as inputs .bedpe or .hic files for 2D contacts and .bedgraph or .bigwig for 1D signals. As these are the widely used, standard format adopted by the NIH ENCODE (ENCODE project consortium, 2004) and 4DN (Reiff et al., 2022) consortia, users do not need to process the data themselves. Not requiring a peak file is also an advantage as some data processing pipelines may not generate them and peak calling also relies on accurate modeling of restriction enzyme cut sites and other protocol-specific considerations (Lareau et al., 2018). Biologically, HiChIA-Rep allows users to specify a genomic region of interest to compare two datasets for potential integration with transcriptome and epigenome profiles, as we have demonstrated for correlating differential genes with loops in MCF10A and MCF7 cell lines (**Figure 4**). We also showed—through ablation experiments—that both the 1D coverage and 2D contacts are needed to accurately quantify the similarity (**Figure 3c-d**).

More broadly, while analyzing PLAC-seq data (**Figure 5**), we pondered upon the ground truth in developmental trajectory of the cells and whether gene transcriptional landscape should define the cell identity. There is no one-to-one correspondence between genome organization and gene transcription, as exemplified in multiple studies. Rapidly depleting protein factors altered or eliminated chromatin loops, yet transcriptional landscapes remained largely unchanged (Rao et al., 2017; Hsieh et al., 2022; Haarhuis et al., 2017); unchanged genes were often connected by short, cohesin-independent loops that were missed in Hi-C data but captured in RNAPII ChIA-PET (Kim et al., 2026). Studies that inhibited transcription showed little change in genome structure (Jiang et al., 2020). Indeed, genome structure and gene transcription are not perfectly correlated, and understandably so: each modality offers fundamentally different information about the cell, showing distinct slices of molecular processes in developmental landscapes. Finding how chromatin interactions affect and are affected by gene transcription in each cell is important in understanding cell fate decisions, which more broadly will help establish necessary or sufficient sources of information when it comes to defining cell identity. These objectives will likely require careful analyses of single-cell multi-omics datasets.

As the 3D genomics field continues to develop methods to jointly profile multiple modalities across a wide range of biological systems, researchers will also need sophisticated algorithms and computational frameworks that can holistically view, compare, and reason within the various modalities. There are additional challenges with single-cell multi-ome data, such as ChAIR (Chai et al., 2024), which can jointly profile chromatin structure, open chromatin region, and gene transcription in a single cell; similarly, HiRES (Liu et al., 2023), GAGE-seq (Zhou et al., 2024), LiMCA (Wu et al., 2024), MUSIC (Wen et al., 2024) simultaneously probe genome folding structure and gene transcription; scHiCAR (Wei et al., 2026) and CHARM (Chen et al., 2026) measure three or more modalities. Indeed, the methods used in HiChIA-Rep can be extended to such data under the computational framework of *structured data* (e.g., graphs). We expect HiChIA-Rep to provide important first steps in jointly leveraging chromatin structure and enrichment signals in multi-modal data, enabling algorithmic and biological insights into capturing and understanding the workings of genome structure and cell function.

## DECLARATIONS

### Availability of data and materials

The datasets supporting the conclusions of this article are included within the article and **Additional File 1**.

The HiChIA-Rep software is available under the MIT License: https://github.com/minjikimlab/hichiarep.

### Authors’ contributions

M.K. conceptualized the project. H.B.Z. wrote the initial ChIA-Rep software package, providing the infrastructure to efficiently pre-process the bedgraph and bedpe files. S.K. devised the graph signal processing algorithms and wrote the HiChIA-Rep Python package with minimal guidance from M.K. J.T.J. contributed scripts to simulate downsampling and tested AREDCI and HPRep. M.K. and S.K. designed tests, interpreted results, and wrote the manuscript. All authors read and approved the final manuscript.

## Acknowledgements

The initial pilot phase of this project was supported by Yijun Ruan at the Jackson Laboratory, where M.K. and H.B.Z. distributed ChIA-Rep software, which is substantially different from HiChIA-Rep. The authors thank: Chia-Lin Wei, Harianto Tjong, Haoxi Chai, Yarui Diao, Xiaolin Wei, and Yueyuan Xu for sharing processed files of ChIATAC and HiCAR data; the ENCODE consortium Nuclear Architecture Working Group members for insightful comments and feedback on the initial ChIA-Rep method; and members of Minji Kim’s research group for helpful discussions.

## METHODS

### Notation

Let the contact matrix of the chromatin interaction frequencies be denoted as *A* ∈ ℝ*^N^*^×*N*^, where *ij*th entry *A_i_*_,j_ encodes the interaction frequency between genomic loci *i* and *j*. The genomic loci are binned to a user-defined resolution (see **Parameters**). Depending on the context, *A* can refer to the entire chromosomal contact matrix or a specific window (see **Sliding Window**), which will be made clear in the section. The indices are 1-indexed and ‘:’ denotes a list of numbers (e.g., 1:5 is the list of numbers 1, 2, 3, 4, 5). Let *b* ∈ ℝ*^N^* denote the corresponding coverage vector, where *b_i_* is the read coverage at genomic locus *i* that is binned to the same resolution as the contact matrix. The coverage vector *b* may either refer to the binding affinity track of a protein or DNA accessibility, depending on whether the data is generated from ChIA-PET (also HiChIP, PLAC-seq) or ChIATAC (also HiCAR), respectively. HiChIA-Rep handles both cases identically and the vector *b* is simply referred to as the ‘coverage vector’ as a placeholder for both situations. Similarly to the contact matrix, *b* may refer to either the entire chromosome or a specific window, which is made clear in the context it is defined.

### Inputs

The main input data to the HiChIA-Rep program are:

1. Contact matrix file (either .hic or .bedpe) to construct *A*.
2. Coverage vector file (either .bedGraph or .bigWig) to construct *b*.
3. UCSC chromosome sizes file to know when to stop the sliding window (see **Sliding Window**).

There are also a few meta-data files, which specify the path locations of the data and which combinations of the data the user wishes to compare. Please see the Github page (https://github.com/minjikimlab/hichiarep) for how to generate these files. HiChIA-Rep can load in multiple samples at once to compare various combinations of pairs efficiently.

### Parameters

All results in this manuscript are generated using the following default parameter settings and window-weighted scores (**Computing Similarity**) with v0.3.0 unless specified otherwise.

The main parameters are the following:

- *window-size* (related to **Sliding Window**), which is the size in basepairs of the sliding window (default: 5,000,000).
- *bin-size,* which determines the size of the bin in basepairs for the contact matrix *A* and the coverage vector *b* (default: 10,000).
- *window-stride* (related to **Sliding Window**), which sets the stride factor for the sliding window (default: 2).
- *mu* (referred to as *μ* in **Methods**) (related to **Signal-Graph Information Exchange**), which is the number of steps in random walks to perform (default: 5).
- *Compare method* (related to **Computing Similarity**), which specifies how to compute the similarity between processed coverage vectors (default: “spearman”).

Additional parameters are the following:

- *chroms_to_load*, which specifies chromosomes to compare (default: “all”).
- *genomic-location*, which specifies a single genomic region “chrom:start-end” (bp) to compare as opposed to a sliding window with option “all” (default: “all”).
- *shapley*, which specifies whether to use ablations to compute window scores comparing contact matrix only or coverage vector only (default: False).
- *output-dir*, which specifies the results folder name (default: “output”).
- *ba-mult*, which is the value to multiply the coverage vector values (default: 1).
- *num-cores*, which specifies the number of cores for parallel processing (default: 1).
- *min-hic-value*, which is the minimum Hi-C interaction value to consider as a valid interaction (default: 1).
- *min-bedgraph-value*, which is the minimum coverage vector value to consider as a valid interaction (default: 1).
- *diagnostic-plots* specifies whether to save diagnostic plots of coverage vector (default: True).
- *do-output-graph* which specifies whether to output .npy graphs for chr1 windows (default: False).

### Sliding Window or User Specified Window

HiChIA-Rep performs a sliding window across the main diagonal of each chromosomal contact matrix *A* ∈ *R^N^*^×*N*^that is binned at the user-specified resolution (**Figure 1d**). The motivation for using a sliding window is that entries near the main diagonal contain most of the biologically relevant structures in 3C data, such as TADs, loops, and stripes and is a common assumption for many Hi-C analysis software (e.g., HiCRep). Since enrichment-based assays select for interactions with proteins or open regions, the structures we observe are typically specific, high-resolution and focused along the main diagonal, making a sliding window method all the more suitable for enrichment-based data as long as the window size is set appropriately.

Specifically, we extract the values of a square region whose size is specified by parameter *window size*. The first sliding window is positioned such that the top left corner of the window aligns with the top left corner of the chromosomal contact matrix. Subsequent windows slide ‘along the main diagonal’, meaning that they are positioned such that the diagonal elements align with the main diagonal of the chromosomal contact matrix. This will ensure that each sliding window is symmetric and interpretable as a small graph (needed for **Signal-Graph Information Exchange**). As of the locations along the main diagonal, we slide each window such that there is a 1/(*windowstride*) overlap between adjacent windows (**Figure 1d** shows the gray and black window for window stride being 2 i.e. ½ overlap). Although adjacent windows may contain the same entries, we implement the window stride feature to ensure that structures which fall between windows are not entirely missed. The sliding window is performed on each chromosome separately until the end of the chromosome, where the last sliding window may end up being smaller than the window size if the chromosome size is not a perfect multiple of the window size. Since many matrix norms and distances are dependent on the dimensions of the matrix, we wanted to ensure that all windows are of the same size. To guarantee this, we position the last window such that the lower right corner coincides with the lower right corner of the chromosomal contact matrix (**Figure 1d**). The sliding window is done similarly for the coverage vector, where we extract entries coinciding with the same genomic regions as the contact matrix (**Figure 1d**, orange vector).

#### Example

- Chromosome size: 23 Mb.
- Window size: 10 Mb.
- Bin size: 10 kb.
- Window stride: 2.
- The contact matrix *A* is dimension *N* × *N*, where *N* = 2300 (23 Mb / 10 kb = 2300).
- The coverage vector *b* is dimension *N* × 1.
- The binned window size is 1000 (10 Mb / 10 kb = 1000).
- Window 1 (genomic location 0-10 Mb)
  - Contact matrix: *A*_1:1000,1:1000_.
  - Coverage vector: *b*_1:1000_.
- Window 2 (genomic location 5-15 Mb)
  - Contact matrix: *A*_500:1500,500:1500_.
  - Coverage vector: *b*_500:1500_.
- Window 3 (genomic location 10-20 Mb)
  - Contact matrix: *A*_1000:2000,1000:2000_.
  - Coverage vector: *b*_1000:2000_.
- Window 4 (genomic location 13-23 Mb)
  - Contact matrix: *A*_1300:2300,1300:2300_.
  - Coverage vector: *b*_1300:2300_.
- Notably, the last window (window 4) is not smaller than the other windows.

Alternative to the sliding window is the option to compare just a single window by providing explicit genomic coordinates (e.g., “chr3:50-647832”). This allows users to compare specific genomic regions of interest without having to compare windows genome-wide. For notation, we assume hereon that we have a sliding window set-up with multiple windows indexed by *k* ∈ {1,2, …}; then, the case of providing genomic coordinates is the special example of *k* ∈ {1} (only 1 window).

#### Unmappable Regions

Hi-C contains entire rows (and columns) of zeros, often near unmappable regions at centromeres or telomeres. Since we divide each row by its sum in *Random Walk Kernel Construction*, we remove the rows (and columns) which sum to zero by keeping the intersection of non-zero rows across the two windows being compared to ensure dimensions are the same. If the number of rows (and columns) kept is less than 10, then we skip computing the window score for this region.

### Signal-Graph Information Exchange

The inputs to this module are a contact matrix and corresponding coverage vector of an arbitrary sliding window (**Figure 1d**, bottom). Since this module is independent of the chromosome or the location along the chromosome of the sliding window, for notational simplicity, let the contact matrix be denoted as *A* ∈ ℝ*^M^*^×*M*^and the corresponding coverage vector *b* ∈ ℝ*^M^*, where *M* < *N* is the number of bins of an arbitrary window.

#### Coverage Weighting

As motivated in the **Results**, coverage weighting is used to normalize the contact matrices analogous to Hi-C normalization methods. Since all pairwise fragments are captured in a Hi-C or Micro-C experiment (**Figure 1a**), the resulting contact matrices are expected to have uniform coverage (i.e. their rows and columns should all sum to the same value). Hi-C normalization (e.g., ICE or KR) are computational methods to address the issue where the actual data may not reflect this invariant, which is problematic if certain parts of the genome are under-represented due to GC bias or technical artefacts. Normalization thus serves the purpose of “bringing out” interactions that are artificially under-represented, enabling downstream algorithms, such as TAD or loop callers to assume that the identified structures are true biological structures, not technical artefacts. ChIA-PET and related enrichment-based technologies require similar preprocessing steps.

A common preprocessing step for ChIA-PET is ‘peak support filtering’, where only loops whose anchors correspond with high coverage vector are kept (**Figure S1a** and **Results**). Operationally, this consists of only keeping interaction *A_i_*_,j_ if *b_i_* and/or *b*_j_ is sufficiently high, where “sufficiently high” is determined by a peak-calling software. This simple operation helps bring out relevant interactions whilst reducing noise (**Figure S1a**). However, the drawbacks include (1) a reliance on the accuracy of peak-calling software and (2) the discrete nature of keeping or filtering interactions based on an arbitrary cutoff threshold, such as determining whether to keep interaction *A_i_*_,j_ if *b_i_* is high AND *b*_j_ is high, or alternatively the OR of the conditions. Importantly, we expect to have greater confidence in interactions *A_i_*_,j_ where *b_i_* is high AND *b*_j_ is high over interactions *A_i_*_j_ where *b_i_* is high XOR *b*_j_ is high. Nevertheless, peak support filtering offers no clear options for how much the former should be emphasized over the latter and is, in general, variable based on the parameters and accuracy of the peak-calling software.

Our coverage weighting scheme is designed to improve upon ‘peak support filtering’ in all the aforementioned points: to not rely on peak-calling software and to emphasize interactions in a continuous manner, prioritizing interactions where both anchor signals are high over cases where only one anchor signal is high. We achieve this using the continuous coverage vector value itself. First, for every genomic loci pair *i*, *j*, we extract the coverage vector values *b_i_*, *b*_j_ and add them, emphasizing interactions between genomic loci *i* and *j* that are well-supported by the coverage vector. This term is then multiplied by the existing loop weight *A_i_*_,j_ to enforce the necessary condition that there must be a positive loop weight for amplification. (**Figure S1b**). In summary, coverage weighting normalization is,

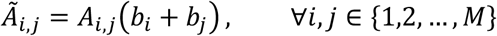

or equivalently,

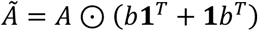

where *Ã* is the coverage weighted contact matrix, ⊙ is the Hadamard product, and **1** is a vector of ones. Coverage weighting ensures that interactions that are well-supported by the enrichment data are emphasized appropriately. At a high level, we are incorporating information from the coverage vector *b* into the contact matrix *A*, thereby reducing noise and technical artefacts in the chromatin contact matrix (**Figure S1c**). Finally, we perform 2D mean filter blurring on *Ã* with fixed filter size of 3 bins, which helps further reduce noise that are particularly present at high resolutions.

#### Random Walk Kernel Construction

The next step is to construct a random walk kernel using the normalized contact matrix from the previous section. A subtle conceptual point is that the matrix *Ã* is viewed as a weighted adjacency matrix associated to a graph with each node corresponding to a binned genomic region. Since *A* now corresponds to a graph, we can apply graph signal processing techniques. If the coverage vector represents protein binding, then random walk can simulate how the proteins might randomly disperse on the DNA through time, jumping randomly but preferentially to regions that are close in 3D nuclear space (**Figure S2d**).

To do this, we first construct a kernel *K*(*μ*) that specifies a random walk with *μ* steps, where *μ* assumes the role of the time parameter. Mathematically, the kernel is defined as,

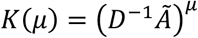

where *D* = *diag* NC*Ã***1**E, …, C*Ã***1**E O is a diagonal matrix containing the row sums and **1** ∈ ℝ^M^ is the all-ones vector. Multiplying *Ã* with *D*^−1^ has the effect of normalizing the rows of *Ã* to sum to 1, making C*D*^−1^*Ã*E*^μ^* a µ-step transition matrix associated to a Markov Chain.

#### Normalization

After the kernel *K*(*μ*) is constructed, we then normalize the coverage vector *b* ∈ ℝ*^M^* such that it sums to 1. Since the coverage vector values are assumed to be non-negative, the normalized vector is a probability mass function. We define the normalized signal as,

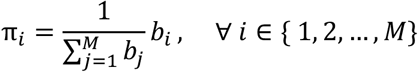

where π ∈ Δ*_M_*_–1_is the normalized coverage vector and Δ*_M_*_–1_is the (*M* − 1)-dimensional probability simplex. The reasoning behind this step is that we care about the strength of the signal relative to other parts of the vector, rather than the absolute value of the signal. Additionally, if we perform random walk on a probability distribution, then the output is also a probability distribution, enabling us to compare the signals using methods developed from information theory (see **Computing Similarity**). For brevity, we will refer to *π* as simply the (normalized) coverage vector.

#### Graph Signal Processing

The graph signal processing step uses the constructed kernel *K*(*μ*) to perform a µ-step random walk of the coverage vector *π*. Mathematically, this corresponds to a single matrix-vector multiplication,

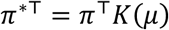

where *π*^∗^ is the coverage vector after performing a *μ*-step random walk. If *π* represented protein binding, then *π*^∗^ describes how the protein would have dispersed through the graph according to the chromatin structure after each protein takes µ random steps. For an example of this effect, see **Figure 2g** and **Figure S2d**.

At a high level, graph signal processing incorporates structural information about the chromatin graph back into the coverage vector, such that *π*^∗^ is a blend of both (1) the normalized coverage vector π and (2) chromatin structure from *A*. We refer to π^∗^ as the ‘processed coverage vector.’

### Computing Similarity

#### Similarity scores for each window

Given two processed coverage vectors corresponding a genomic window, one from input sample 1 and another from input sample 2, we compute the window score using either the Spearman correlation or a transformed Jensen-Shannon Divergence (JSD) (Lin et al., 1991) (**Figure 1d**). Whether to use the Spearman correlation or JSD is a user-defined parameter.

Let the processed coverage vectors be *π^A^* ∈ Δ*_M_*_–1_ and *π^B^* ∈ Δ*_M_*_–1_, where *M* is the number of bins of an arbitrary window. The window score *ρ_c_*_,*k*_ for the *k*th window of the sliding window and *c*th chromosme is defined as

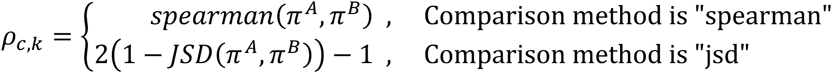

where we simply take the Spearman correlation between the vectors *π^A^* and *π^B^* if compare method parameter is “spearman” or a transformed JSD measure if “jsd”. The *JSD* is the Jensen-Shannon Divergence and is

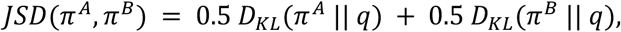

where

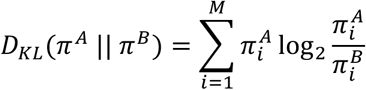

is the KL divergence (Kullback & Leibler, 1951) between distributions *π^A^* and *π^B^* and

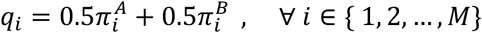

is the mixture distribution.

The transformation after computing the JSD is to convert the divergence into a similarity score in the range [−1,1], where a high value indicates high similarity.

#### Aggregating similarity scores for each window

Let the number of sliding windows for the *c*th chromosome be *K_c_*. Assume that we compute the window score across the sliding window, obtaining a sequence of window scores *ρ_c_*_,1_, *ρ_c_*_,2_, …, *ρ_c_*_,*K*_*_c_* for chromosome *c* ∈ {1,2, … *C*}. Then, the chromosomal reproducibility score *ρ_c_* between input sample 1 and input sample 2 is the average of the window scores in the chromosome:

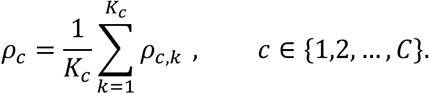

The unweighted genome-wide HiChIA-Rep score *ρ_HiC_*_ℎ*IA*–*Rep*_ between input sample 1 and input sample 2 is the average of the chromosomal reproducibility scores *ρ*_1_, *ρ*_2_, …, *ρ_C_* i.e.,

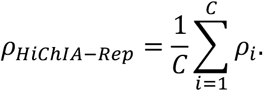

In the **Sliding Window or User Specified Window** section, we removed a fixed set of rows (and columns) which summed to 0. Denote the fraction of rows kept for the *k*th window comparison of the *c*th chromosome be *β_c_*_,*k*_. We then construct a probability mass function of normalized window weights for the chromosome: for each chromosome *c* ∈ {1,2, …, *C*}, define window weights

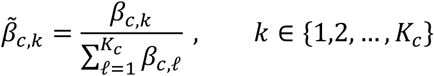

such that we compute the window-weighted chromosomal reproducible score *ρ̃_c_*:

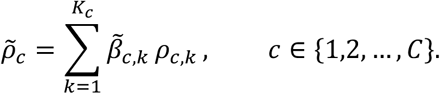

The interpretation is that we weigh window score *ρ_c_*_,*k*_ lower, relative to other windows in the chromosome, if many rows have been removed in window *k*, for reasons such as being near unmappable regions. Since there are fewer genomic loci being compared in these cases, the weighted average is designed to downweigh these window scores according to their expected contribution. Then, the window-weighted genome-wide HiChIA-Rep score is,

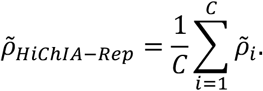

### Ablation tests

We performed ablations on input data to quantify the respective contributions of contact matrix *A* and coverage vector *b* to the reproducibility score. We considered three choices for ablation tests: *identity elements*, input control or the Hi-C expected profile, and shuffled inputs.

#### Ablation Type 1: Identity Elements

For the identity element ablations, we either replaced (i) the contact matrix *A* with the identity matrix *I* to ablate contact matrix, or (ii) the coverage vector *b* with the all-ones vector 1 to ablate coverage vector. The motivation for these choices is that the identity matrix is the identity element for matrix-vector multiplication, which is the operation used in the *μ*-step random walk (*Graph Signal Processing*), and the all-ones vector is the identity element for the Hadamard product, which is used in coverage weighting (*Coverage Weighting*).

In the following, we trace through the effect of each ablation.

(i) **Ablate contact matrix:** *A* = *I*. If *A* = *I*, the coverage weighting step becomes

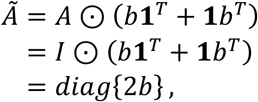

yielding a diagonal matrix. After the 3×3 blur, we obtain a matrix that is approximately diagonal with positive entries restricted to strata (i.e., off-diagonals) 0, 1, 2 and everywhere else being filled with zeros. The next step is *Random Walk Kernel Construction*, where we divide each row by the row sum of *A* so that each row sums to 1. This means that the random walk kernel will approximate an identity matrix, and the effect of graph signal processing on the coverage vector *π* is a blurring operation, effectively removing information in the contact matrix and only taking into consideration the coverage vector.
(ii) **Ablate coverage vector:** *b* = **1**. If *b* = **1**, the coverage weighting becomes

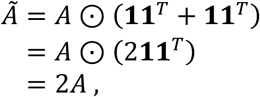

yielding a scalar multiplication of the contact matrix *A*. The 3×3 blur and the row sum normalization in *Random Walk Kernel Construction* will mean that the kernel is a blurred contact matrix *A* with row normalized to sum to 1. In the graph signal processing section, the coverage vector *π* will be

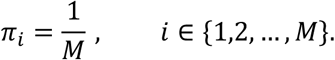

Thus, the effect of the *μ*-step random walk is interpretable as an interpolation from uniform distribution to the stationary distribution. In any case, coverage vector information has been removed from influencing the scores.

For each window *k* ∈ {1,2, …, *K_c_*} on chromosome *c* ∈ {1,2, …, *C*}, we compute the following similarity scores:

- Window score (unablated): *ρ_c_*_,*k*_.
- Window score of coverage vectors: ρ^*V*^_*c,k*_, where ‘V’ means that the coverage vector is left intact and we ablate the contact matrix with the identity matrix substitution *A* = *I*.
- Window score of contact matrices: ρ^*M*^_*c,k*_, where ‘M’ means that the contact matrix is left intact and we ablate the coverage vector with the all-ones substitution *b* = **1**.

If *shapley* parameter is True, then the program reports all three scores, whereas if *shapley* is False then only the unablated window score is reported.

#### Ablation Type 2: Input Control and Expected Profile

The input control is a baseline reference for IP-based coverage tracks (e.g., ChIP-seq), designed to capture the randomness and biases in the experimental protocol, such that significance of observed peaks can be evaluated with respect to this baseline. While a direct counterpart for chromatin interaction data does not exist, the Hi-C expected profile, which is the mean of each stratum, is used for a similar purpose: as a relative baseline to infer the significance of observed frequencies (O/E computation). Thus, the input control and expected profile are potential ablation input data that are closer to the data distribution than identity elements. For instance, the expected profile captures the distance-dependent decay of chromatin interaction data, whereas the identity matrix does not. We therefore used input control and the Hi-C expected profile as another type of ablation.

For the coverage vector ablation, we replaced the observed coverage vector with input control of GM12878 ChIA-PET. For the contact matrix ablation, we replaced the observed contact matrix with the Hi-C expected profile of GM12878 Hi-C data (4DNFI1UEG1HD).

#### Ablation Type 3: Shuffled Locations

The third ablation type aims to ensure the ablate input data is as close to the distribution of data as possible. Here, we shuffled the genomic locations –for either the contact matrix or coverage vector– to remove genomic correspondence while retaining properties of the observed data. Specifically, for the coverage vector ablation, we shuffled the genomic locations of the coverage vector within a chromosome. Similarly for the contact matrix ablation, we shuffled the genomic locations for the contact matrix. We note that the genomic span (the range from ‘start’ to ‘end’) was kept the same for shuffled locations.

### Computing a reproducibility threshold for HiChIA-Rep

Given HiChIA-Rep non-replicate points {*x*_1_, …, *x_n_*} and replicate points {*y*_1_, …, *y_m_*}, we seek to find a suitable threshold for reproducibility. We were inspired by the hard-margin classifier (Yardimci et al., 2019) that determines the threshold to be equidistant from the boundary of each class, that is, between 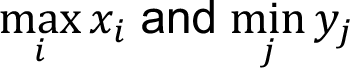. However, the max and min functions are neither robust to outliers and nor do they take into account the *distribution* of the data points. Thus, at a high level, we aim to keep the essence of equidistance between boundary points, but between boundary points that consider the measure of center and spread of the distribution.

For illustrative purposes, let the measure of center be the mean *μ_x_*, *μ_y_* (assume without loss of generality *μ_x_* < *μ_y_*) and the spread be the standard deviation *σ_x_*, *σ_y_*. Specifically, we want the threshold *ε*^∗^ to be equidistant between the two means *μ_x_*, *μ_y_* (i.e., *μ_x_* < *ε*^∗^ < *μ_y_*) in units of standard deviation. Equivalently, we wish to find threshold *ε*^∗^ ∈ [*μ_x_*, *μ_y_*] whose Z-scores with respect to the two groups are equal:

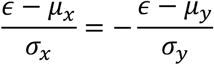

where solving for *ε* yields the solution:

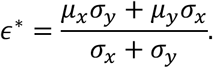

For interpretation, it helps to re-express the solution as a convex combination between the two means *μ_x_*, *μ_y_* i.e.,

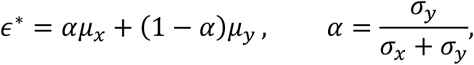

which gives us the fact that the solution *ε*^∗^ lies between the two means *μ_x_* < *ε*^∗^ < *μ_y_*. Here, *α* ∈ [0,1] is 0.5 if the standard deviations are the same *σ_x_* = *σ_y_*. If *α* < 0.5, then *σ_y_* < *σ_x_* and *ε*^∗^ is closer to *μ_y_* than *μ_x_*, which is as we expect.

In practice, since the number of replicate points is small, we used the median as a robust estimate of the center, and the median absolute deviation as the estimate for spread. The non-replicate and replicate points used to determine the threshold is from various cell-lines (HFFc6, MCF7, MCF10A, HEK293T, S2, GM12878, HG00731, H1, B6 mouse), technologies (ChIA-PET, ChIATAC, HiCAR, HiChIP, PLAC-seq) and enrichments (CTCF, RNAPII, SMC1A, RAD21, open regions, H3K4me3), amounting to 465 non-replicate points and 43 replicate points for *ε*^∗^ = 0.894.

### Running AREDCI and HPRep

We modified the ChIA-PET2 pipeline (v0.9.3) (Li G et al., 2017; He et al., 2015) to utilize bedpe ChIA-PET data with PET (paired-end tag) count ≥ 3 to generate MICC files as input to AREDCI (from a commit on Jun 3, 2024) (Sun et al. 2024). Since the bedpe files contain intra-chromosomal PETs, we first converted them into ChIA-PET2’s PET format wherein each cluster was expanded into rows according to the PET count column, skipping steps 1–3 and deduplication (linker trimming, mapping and PET construction). MACS2 peak calling (step 4) was run in BEDPE mode, and MICC (step 7) was run through a wrapper to accept an empty inter-chromosomal file. All other steps were unchanged and used default parameters. For AREDCI, parameters were left as their default except for disabling DCI identification and using the hg38 blacklist file.

We ran HPRep (from a commit on Dec 16, 2021) (Rosen et al. 2021) pipeline with our own scripts on ChIA-PET peak files and bedpe files with PET count ≥ 3. Modification through an adapter was required since HPRep is intended for raw PLAC-Seq or HiChIP data, not ChIA-PET. The adapter closely replicated *split_and_xor.py* using same-bin contacts as a proxy for *chrt.vip.bed* and ran the original pipeline thereafter, starting with *find_anchor_bins.R*. Specifically, the adapter consists of *bedpe_to_binned_pixels.py* and *build_reg_raw.py* which transform a bedpe and peak file to AND and XOR files required by *find_anchor_bins.R*.

First, *bedpe_to_binned_pixels.py* converts raw bedpe contacts into a tsv file containing binned contacts and their corresponding frequency. Our binning convention exactly matches that of *split_and_xor.py* in the original pipeline. Second, *build_reg_raw.py* takes the binned tsv file and peaks file as input and outputs AND and XOR files. The original pipeline’s *chrt.vip.bed* file cannot be generated from our data, so we compute the short count per bin by using same-bin bedpe contacts as a proxy. Peaks are then binned and output files are generated according to *split_and_xor.py* exactly as in the original pipeline: The AND file is constructed from bedpe pairs that intersect a peak at both anchors. The XOR file is constructed from bedpe pairs that intersect a peak at exactly one anchor.

All HPRep parameters were left as default as in *bin/run_pipeline_stages_1_2.sh* and *bin/run_pipeline_stage_3.sh*, except the genomic features file and the resolution of 50kb. The genomic features file was set to AluI rather than MboI since the ChIA-PET protocol detailed on 4DN/ENCODE used AluI as the restriction enzyme. We document all modifications in the github page (https://github.com/minjikimlab/hichiarep).

### Running Hi-C reproducibility methods

We ran the python implementation of HiCRep (v0.2.6) (Yang et al. 2017), and GenomeDISCO, HiC-Spector, and QuASAR-Rep using the combined package 3DChromatin_ReplicateQC (v0.0.1) (Yardımcı et al., 2019).

#### HiCRep

We ran HiCRep (Yang et al. 2017) at 50 kb resolution with *maxdist=5000000* (5 Mb) to match the window size for HiChIA-Rep. The smoothing parameter *h=5* based on the recommendations from Yang et al. 2017 (https://github.com/qunhualilab/hicrep).

#### GenomeDISCO

We ran GenomeDISCO (Ursu et al., 2018) at 50 kb resolution with the following parameter settings: *subsampling*=*lowest, tmin=3, tmax=3, norm=sqrtvc, scoresByStep=no, removeDiag=yes, transition=yes*.

#### HiC-Spector

We ran HiC-Spector (Yan et al., 2017) at 50 kb resolution with *n=20*.

#### QuASAR-Rep

We ran QuASAR-Rep (Sauria and Taylor, 2017) at 50 kb resolution with *rebinning=‘resolution’*.

### Data Processing

#### Converting .bam to .bedgraph

We generated .bedgraph files from .bam files for all the PLAC-seq coverage vector data (see **Additional File 1**) using samtools (v1.21) (Danecek et al., 2021) and bedtools2 (v2.31.1) (Quinlan et al., 2010). Specifically, we ran *samtools sort*, *samtools rmdup* with option *-s*, and *bedtools genomecov* with options *-ibam* and *-bg*.

#### Converting .pairs to .hic

We generated .hic files from .pairs files for all the PLAC-seq chromatin contacts data (see **Additional File 1**) using JuicerTools (v1.22.01) (Durand et al., 2016) *pre* function with default parameters.

### Analysis

#### Hierarchical Clustering

Hierarchical clustering was performed using *seaborn.clustermap* function with parameters *method=“average”, metric=“euclidean”* from the seaborn package (v0.13.2) (Waskom, 2021).

#### Runtime Analysis

Runtime analysis was performed on a HPC environment using 4 core CPU (3.0 GHz Intel Xeon Gold 6154) and 64GB RAM. A local environment with 10 core CPU (1.8 GHz Intel(R) Core(TM) i7-1365U) and 32GB RAM was used specifically for enrichment-based methods due to the graphical user interface of AREDCI.

#### Sequencing Depth Adjustment

We generated subsampled chromatin contact matrices and corresponding coverage vector for HFFc6 ChIA-PET data (**Figure 2c, d**) using a binomial sampling scheme. Let *p* ∈ [0,1] denote the percentage of further downsampling that we wish to perform (e.g., the *p* for 100%, 50%, 25% and 10% in **Figure 2c** is 1.0, 0.5, 0.25, and 0.1, respectively). Given input sample 1 contact matrix *A*^(1)^ ∈ ℝ*^M^*^×*M*^and coverage vector *b*^(1)^ ∈ ℝ*^M^* and input sample 2 contact matrix *A*^(2)^ ∈ ℝ*^M^*^×*M*^and coverage vector *b*^(2)^ ∈ ℝ*^M^*, we perform subsampling twice. The first round of subsampling is to match, in expectation, the higher depth matrix (or vector) to the lower one. The second round of subsampling takes these matrices (or vectors) and then further subsamples both to the specified fraction *p* of reads.

For the first round of subsampling, we count the total sum of the contact matrices, i.e.,

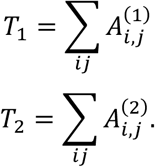

If *T*_1_ = *T*_2_, then the first round of subsampling can be skipped. Otherwise, assume without loss of generality that *T*_1_ > *T*_2_. We subsample the higher depth matrix *A*^(1)^ by sampling reads from a Binomial distribution with success probability *T*_2_/*T*_1_,

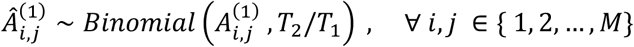

and leave the lower depth matrix unchanged,

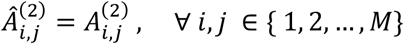

where Â^(1)^ and Â^(2)^ are contact matrices after the first round of subsampling. This procedure is performed similarly for the coverage vectors except for vectors instead of matrices.

The second round performs additional subsampling from a binomial distribution with success probability *p*, i.e.,

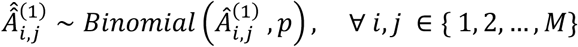

and

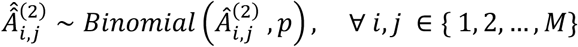

where Â^(1)^ and Â^(2)^ are contact matrices after both round of subsampling. This procedure is performed similarly for the coverage vectors.

#### Chromatin Graph Plot

The chromatin graph plots in **Figure 2/S2** were generated by applying non-metric multi-dimensional scaling (MDS) on the contact matrix *A* ∈ ℝ*^M^*^×*M*^. Specifically, we first extract the coverage weighted contact matrix *Ã* from the respective samples (see **Coverage Weighting**). We then compute a matrix of discrepancies *Y* by

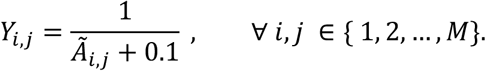

We then used the shortest-paths distances using the *floyd_warshall_numpy* function from NetworkX (v3.4.2) (Hagberg et al., 2007),

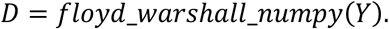

These transformations convert the contact matrix into a matrix of geodesic distances for the MDS algorithm. To obtain the embedding coordinates of each genomic loci, we ran non-metric MDS on *D* using *sklearn.manifold.MDS* with dimension 2 from the scikit-learn package (v1.7.2) (Pedregosa et al., 2011).

#### MCF7 and MCF10A differential gene expression analysis

We downloaded processed DESeq2 results from Barutcu et al., 2015 (GSE71862) for the volcano plot (**Figure 4a**). We classified genes into ‘downregulated’, ‘upregulated’, ‘unchanged’, and ‘other’ (where upregulated means higher in MCF7 compared to MCF10A) by requiring base mean expression ≥ 10, adjusted p-value < 0.1 and log2FC < -1.5 for downregulated, base mean expression ≥ 10, adjusted p-value < 0.1 and log2FC > 1.5 for upregulated, and adjusted p-value ≥ 0.1 and absolute log2FC < 1 for unchanged. All other genes were classified as ‘Other’. We then associated each gene to a window among non-overlapping sliding windows of HiChIA-Rep run at 5 Mb window size, where if any part of the gene body overlaps the window, then it is associated; a gene overlapping two windows was assigned the mean of their scores (**Figure 4c**). In **Figure 4c**, we note that a downregulated gene X and an upregulated gene Y may correspond to the same window score if the two genes X and Y are in the same sliding window, which would weaken the discrepancies between boxplots. For boxplots, we use the two-sided Mann-Whitney U test using *scipy.stats.mannwhitneyu*.

Separately, we downloaded MCF7 and MCF10A gene quantification files from ENCODE (ENCFF721BRA, ENCFF215WRT) to obtain TPM values. We removed gene with TPM less than 5 in both MCF7 and MCF10A and computed the absolute log2FC for each gene using TPM values. We then associated each gene to windows as described above. For each window, we compute the average of the absolute value of log2FC for **Figure 3d** and **Figure S3c**, **d**. We fit scatter points with OLS and performed the two-sided Spearman’s rank correlation test using *scipy.stats.spearmanr* from scipy package (v.1.15.2).

#### MDS on HiChIA-Rep scores

To embed forebrain PLAC-seq samples in 2D coordinates, we used non-metric MDS on the pairwise HiChIA-Rep score matrix. Let the HiChIA-Rep score matrix be denoted as *P*^12*x*12^, where we have 12 samples (FB E12.5, E13.5, E14.5, E15.5, E16, P0 with 2 replicates each). We converted the scores into a dissimilarity matrix i.e.,

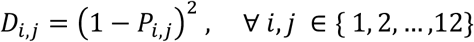

Since the scores in *P* were between 0 and 1, this transformation is monotonic and will preserve the rank order of proximities. We then ran non-metric MDS on *D* using *sklearn.manifold.MDS* with dimension 2 from the scikit-learn package (v1.7.2) (Pedregosa et al., 2011).

#### Genome Browser Plot

Tracks were visualized using BASIC Browser (Lee et al., 2020) and contact matrices were visualized using Juicebox Web App (v2.3.6) (Robinson et al., 2018), if not generated by our custom scripts.

---

**Figure S1:**
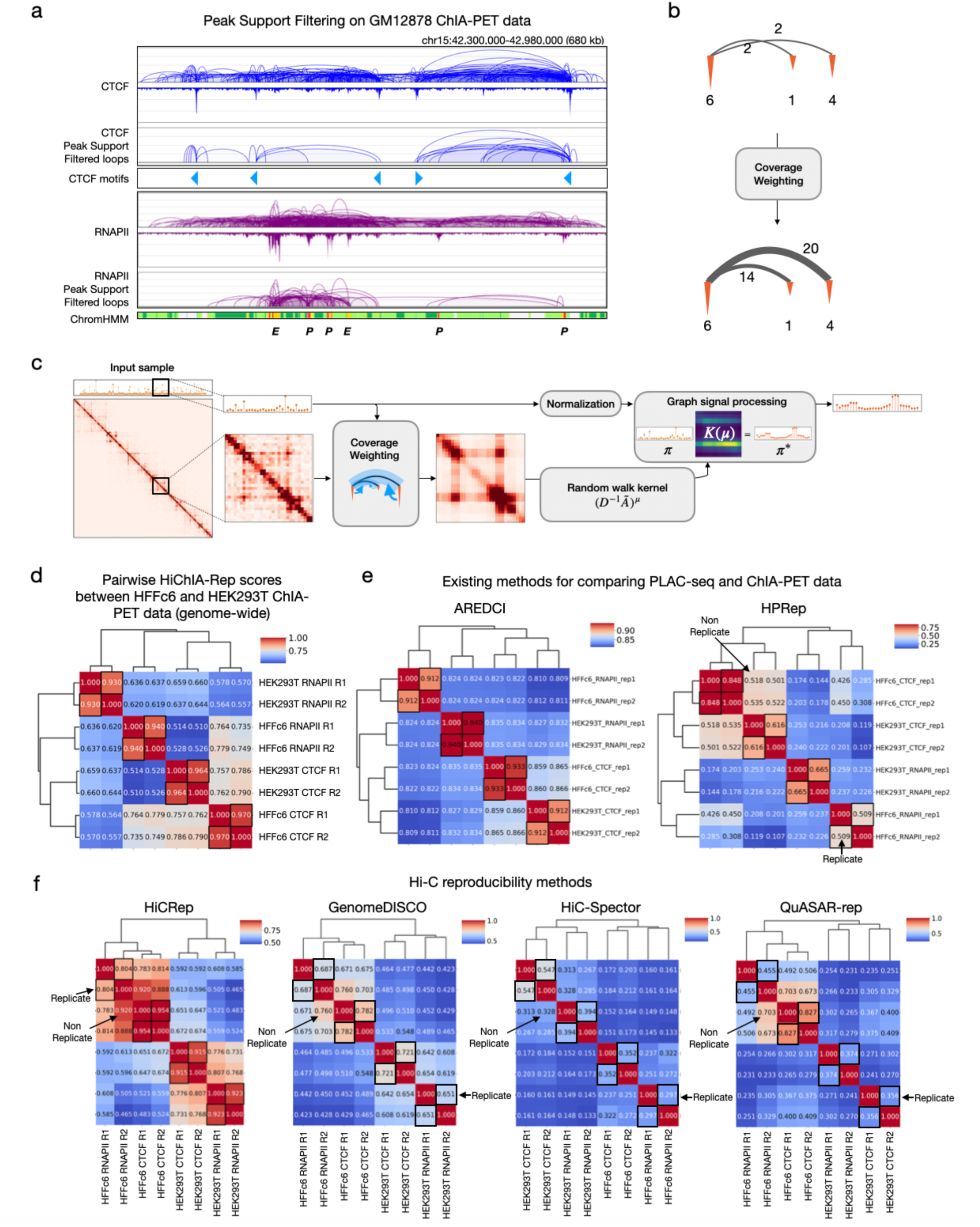
HiChIA-Rep schematics and comparison of methods on HFFc6 and HEK293T ChIA-PET data. Related to **Figure 1**. **(a)** Genome browser tracks showing GM12878 data on chr15:42,300,000-42,980,000: CTCF ChIA-PET ‘loops and coverage’ (row 1), CTCF ChIA-PET loops after filtering for peak support (row 2), CTCF motifs (row 3), RNAPII ChIA-PET ‘loops and coverage’ (row 4), RNAPII ChIA-PET loops after filtering for peak support (row 5), and ChromHMM chromatin states (row 6). Promoters ‘P’ (red), gene transcription (green), enhancers ‘E’ (yellow). **(b)** Schematic of coverage weighting operation on an examplary loops and coverage. The loop weight is the sum of the two anchors’ coverage vector multiplied by the loop weight (e.g. 14 = 2(6 + 1)). **(c)** Signal graph information exchange module on real data: MCF7 CTCF ChIA-PET replicate 1 at chr1:30-33Mb. The inset shows a window chr1:34.75-35.05Mb. The number of steps in random walk is *μ* = 3. **(d)** HiChIA-Rep scores between all pairwise combinations of HFFc6 and HEK293T ChIA-PET data with hierarchical clustering between sample. Replicate pairs are boxed in black. **(e)** Existing reproducibility methods for enrichment-based data (AREDCI and HPRep) on HFFc6 and HEK293T ChIA-PET data. **(f)** Hi-C reproducibility scores (HiCRep, GenomeDISCO, HiC-Spector, QuASAR-rep) on HFFc6 and HEK293T ChIA-PET. Arrows indicate cases in which non-replicate scores are higher than replicate scores.

**Figure S2:**
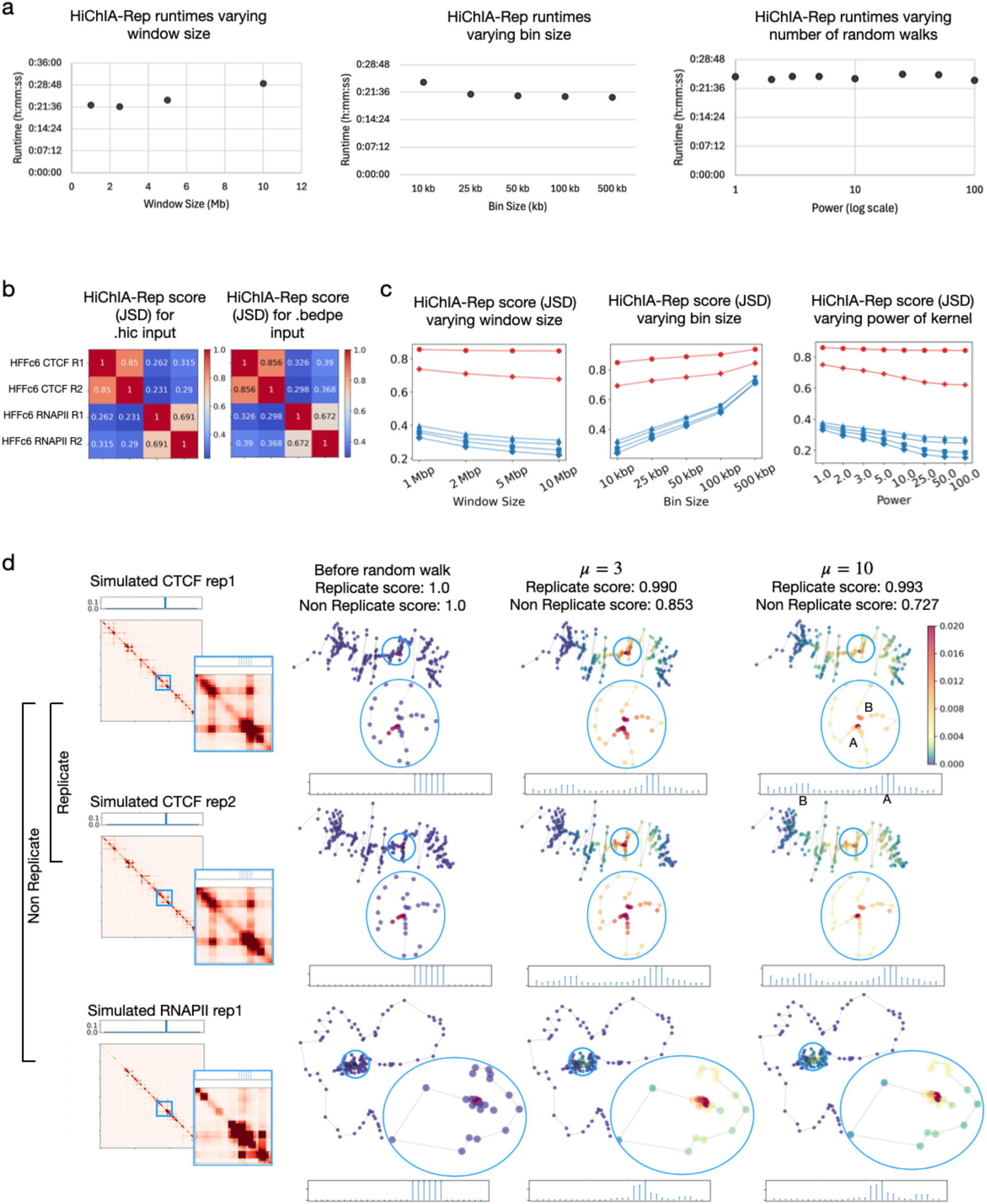
HiChIA-Rep runtimes and effect of varying inputs and parameters. Related to **Figure 2**. **(a)** HiChIA-Rep total runtimes on HFFc6 CTCF (rep1 and rep2) and RNAPII (rep1 and rep2) ChIA-PET data encompassing a total of 6 pairwise combinations, varying window size (left), bin size (center), and number of steps in random walk (right). **(b)** HiChIA-Rep scores using JSD on HFFc6 CTCF and RNAPII ChIA-PET data for .hic input (left) and .bedpe input (right). **(c)** HiChIA-Rep scores using JSD on HFFc6 CTCF and RNAPII ChIA-PET data varying window size (left), bin size (center), and number of steps in random walks (right). **(d)** Left: Contact matrix of MCF7 CTCF (rep1, rep2) and RNAPII rep1 ChIA-PET data on chr1:30-33Mb region. The coverage vector is a delta function placed at chr1:31,920,000-31,980,000. Inset (blue border) shows chr1:31,750,000-32,050,000. Right: MDS embedding of contact matrix with color of points indicating enrichment signal before random walk, after random walk (*μ* = 3), after random walk (*μ* = 10). The scores are the Spearman coefficient of the chr1:30-33Mb window. The coverage vector zoomed into the same inset region is shown below each graph. For the coverage vector shown in *μ* = 10, the loop anchor regions are A and B.

**Figure S3:**
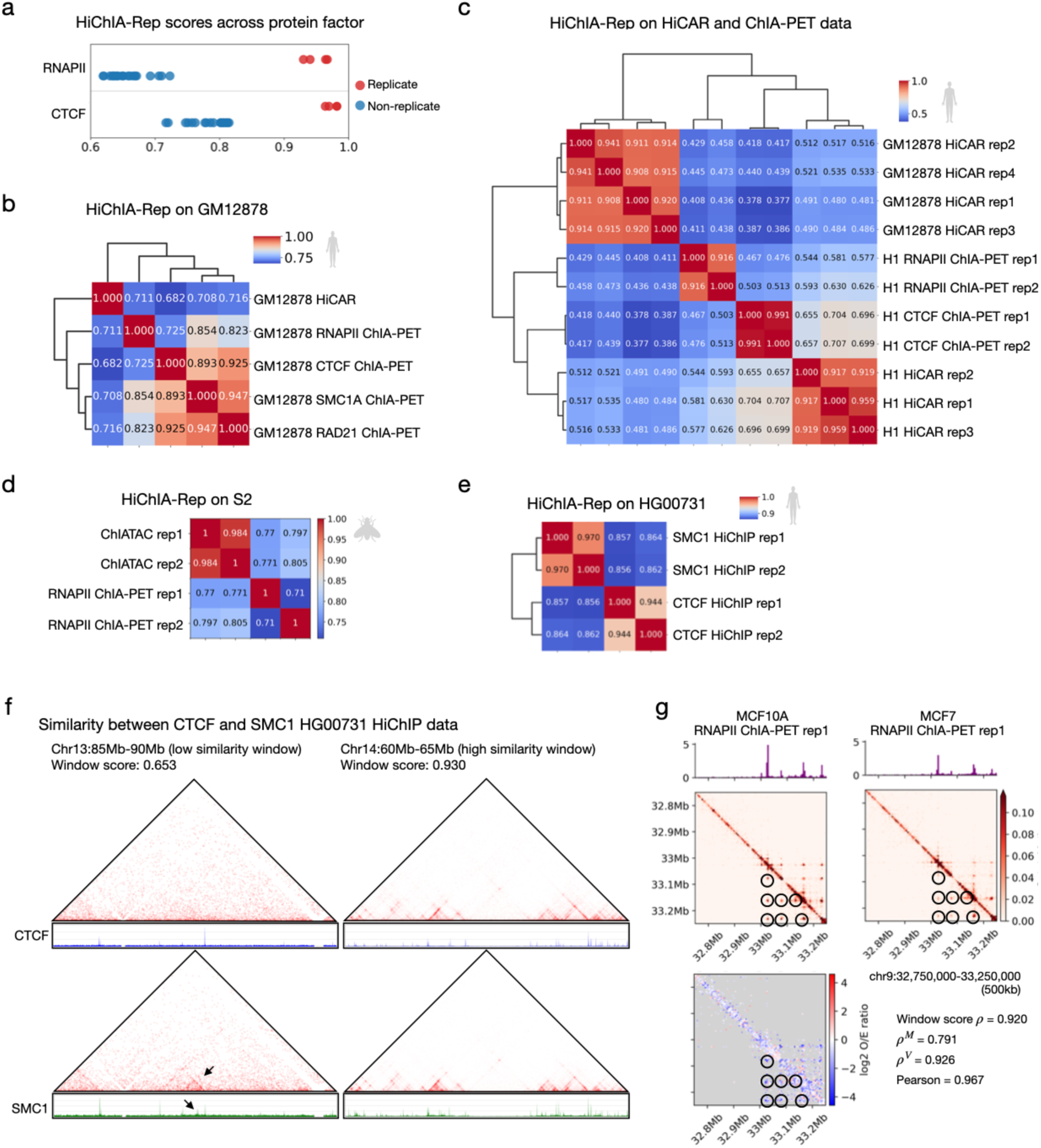
Evaluation of HiChIA-Rep on other cell lines and enrichment-based technologies. Related to **Figure 3**. **(a)** Dot plot of HiChIA-Rep scores on the same dataset as Figure 3a stratified by protein factors RNAPII and CTCF. For instance, in row 1, red points are replicates between RNAPII samples (e.g., HFFc6 RNAPII R1 vs R2) and blue points are non-replicates between RNAPII samples (e.g., HFFc6 RNAPII R1 vs. MFC10A RNAPII R1). **(b)** HiChIA-Rep scores between all pairwise combinations of GM12878 HiCAR (rep1) and ChIA-PET for RNAPII (rep1), SMC1A (rep1), RAD21 (rep1) and CTCF (rep1) with hierarchical clustering. **(c)** HiChIA-Rep scores on GM12878 HiCAR (rep1-4), H1 cell-line ChIA-PET enriched for RNAPII (rep1 and rep2) and CTCF (rep 1 and rep2), and H1 HiCAR (rep1-3) with hierarchical clustering. **(d)** HiChIA-Rep scores on S2 cell line: ChIATAC (rep1 and rep2) and RNAPII ChIA-PET (rep1 and rep2). **(e)** HiChIA-Rep scores on HG00731 cell line: SMC1 HiChIP (rep1 and rep2) and CTCF HiChIP data (rep1 and rep2). **(f)** HG00731 HiChIP CTCF and SMC1 data (replicate 1) Juicebox and genome browser plots showing the contact matrix and coverage vector, respectively, of a 5 Mb region with low HiChIA-Rep window score (left) and high HiChIA-Rep window score (right). Juicebox resolution is 10 kb with normalization “None”. **(g)** An example region on chr9 showing window scores comparing MCF10A and MCF7 RNAPII ChIA-PET: unmodified *ρ*, window score of contact matrix only *ρ^M^*, window score of coverage vector only *ρ^V^* and Pearson correlation coefficient between coverage vectors. Bottom left: Log2 of ratio between MCF7 and MCF10A O/E (observed/expected) normalized contact matrices.

**Figure S4:**
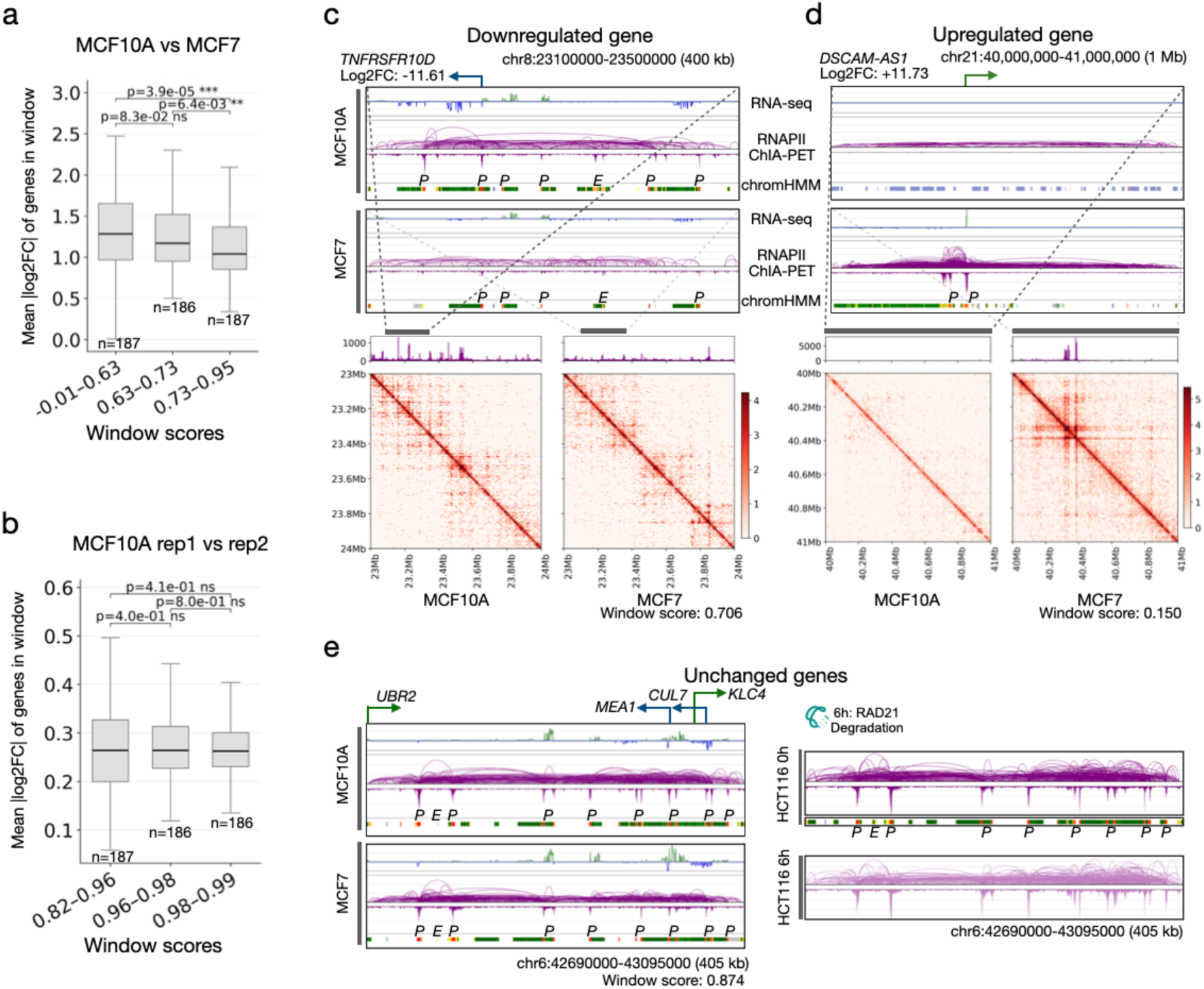
Additional statistical analysis on differential gene expression and HiChIA-Rep scores. Related to **Figure 4**. **(a)** Boxplot of HiChIA-Rep window scores comparing MCF10A and MCF7 RNAPII ChIA-PET data, grouped into approximately equal number buckets ‘-0.01-0.63’ (n=187), ‘0.63-0.73’ (n=186), and ‘0.73-0.95’ (n=187) against the mean absolute log2FC of expressed genes in each window. P-values are from the two-sided Mann-Whitney U test. **(b)** Similar to panel **(a)** but using HiChIA-Rep window scores comparing MCF10A rep1 and rep2 RNAPII ChIA-PET data. **(c)** Example of a downregulated gene *TNFRSF10D* in MCF7 with browser plots of RNA-seq, ChIA-PET ‘loops and coverage’, and chromHMM states (‘P’ promoter in red, ‘E’ enhancer in yellow) for MCF10A and MCF7 (top). Bottom: Coverage vector and contact matrix of a larger genomic range encompassing the *TNFRSF10D* gene with HiChIA-Rep window similarity score. **(d)** Similar to panel **(c)** but showing an upregulated gene *DSCAM-AS1*. **(e)** Left: Browser visualization of select unchanged genes on chr6, showing RNA-seq, ChIA-PET ‘loops and coverage view’ and chromHMM states for MCF10A and MCF7 with window score from HiChIA-Rep program. Right: Browser plot of same region showing ChIA-PET and chromHMM states for HCT116 0hr and 6hr RAD21 depletion. 0hr: no auxin, 6hr: auxin treatment.

**Figure S5:**
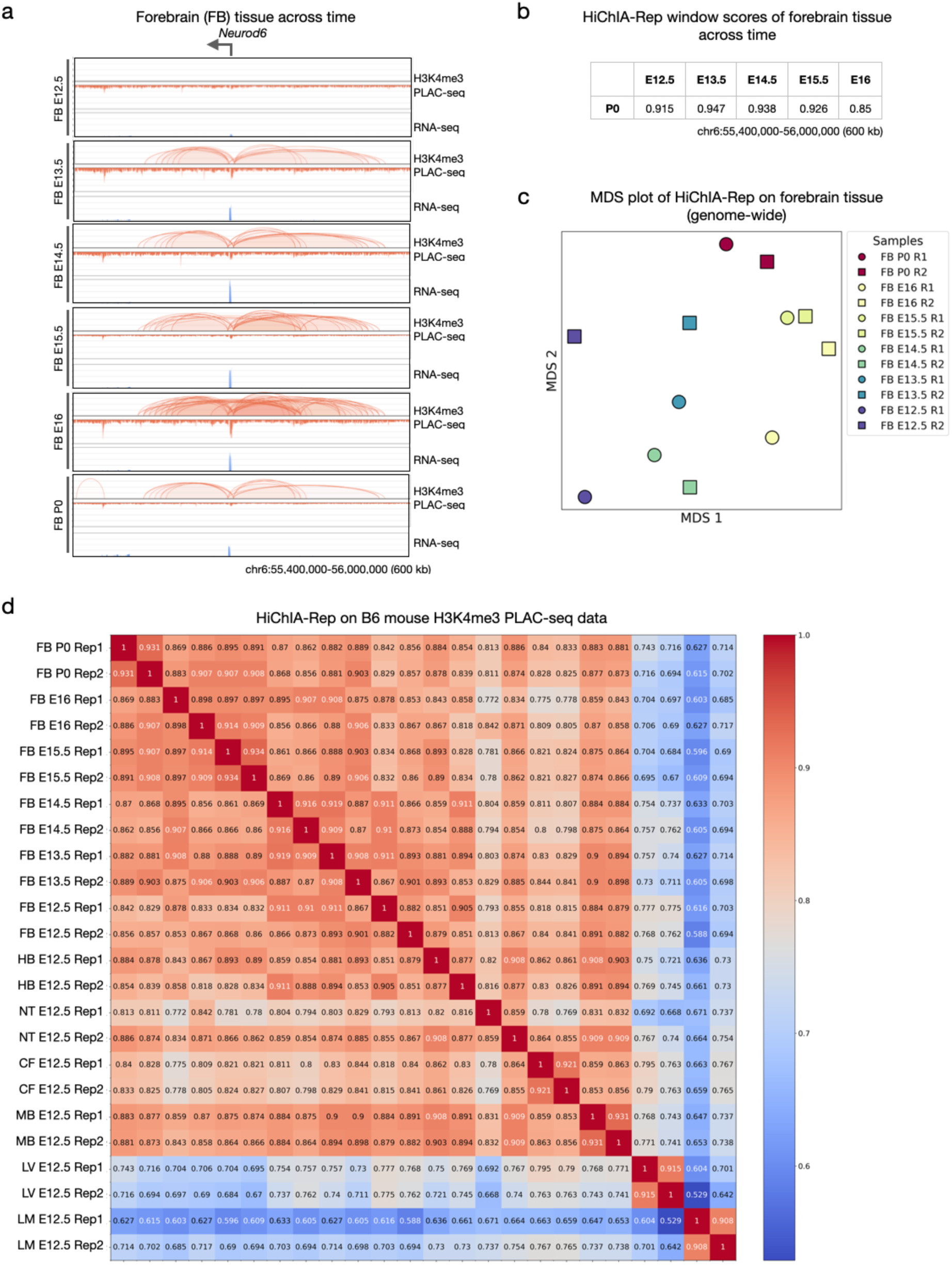
HiChIA-Rep on PLAC-seq data across time. Related to **Figure 5**. **(a)** Browser tracks on mouse forebrain (FB) tissue (chr6:55,400,000-56,000,000) across time: PLAC-seq ‘loops and coverage’ and RNA-seq for forebrain tissue at E12.5 (row 1), E13.5 (row2), E14.5 (row 3), E15.5 (row 4), E16 (row 5), P0 (row 6). **(b)** HiChIA-Rep window scores of the data presented in panel **(a)**. **(c)** MDS (multidimensional scaling) coordinates using genome-wide HiChIA-Rep scores on mouse forebrain tissue. **(d)** HiChIA-Rep scores computed on the entire mouse embryo PLAC-seq dataset, consisting of various tissues (forebrain, hindbrain, neural tube, craniofacial prominence, midbrain, liver, limb) at distinct time points (E12.5-E16, P0). FB: forebrain, HB: hindbrain, NT: neural tube, CF: craniofacial prominence, MB: midbrain, LV: liver, LM: limb, Rep: replicate.

